# The mammalian rapid tRNA decay pathway is critical for N⁷-methylguanosine-hypomodified tRNA degradation under physiological conditions

**DOI:** 10.1101/2025.11.05.686800

**Authors:** Keita Miyoshi, Kiito Otsubo, Shunya Kaneko, Makoto Terauchi, Yuki Hatoyama, Hideki Noguchi, Masato T. Kanemaki, Kuniaki Saito

**Affiliations:** Invertebrate Genetics Laboratory, Department of Chromosome Science, National Institute of Genetics, Research Organization of Information and Systems (ROIS), Shizuoka, Japan; Graduate Institute for Advanced Studies, SOKENDAI, Shizuoka, Japan; Center for Genome Informatics, Joint Support-Center for Data Science Research, Research Organization of Information and Systems (ROIS), Shizuoka, Japan; Molecular Cell Engineering Laboratory, Department of Chromosome Science, National Institute of Genetics, Research Organization of Information and Systems (ROIS), Shizuoka, Japan; Department of Biological Science, Graduate School of Science, The University of Tokyo, Tokyo, Japan

**Keywords:** rapid tRNA decay pathway, XRN2, m^7^G, human HCT116 cell line, physiological conditions

## Abstract

Chemical modifications of transfer RNAs (tRNAs) are integral to their stability and to translation. Loss of N^7^-methylguanosine (m^7^G) on specific tRNAs reduces their steady-state abundance and impairs translation in mammals, but whether these decreases reflect active degradation under physiological growth conditions is unresolved. Here, using human HCT116 cells, we show that knockdown of the 5′→3′ exonuclease, XRN2, restores tRNA levels diminished by METTL1 depletion. Leveraging conditional protein knockdown, we performed time-resolved measurements of mature tRNA levels and directly quantified decay kinetics. We show that in the absence of heat stress, m⁷G-hypomodified tRNAs undergo XRN2-dependent accelerated decay. Finally, partial loss of the *Drosophila* XRN2 ortholog, Rat1, genetically rescues male sterility of *mettl1* mutants, demonstrating organismal relevance. These findings define a conserved constitutive rapid tRNA decay pathway in mammals and indicate inhibition of tRNA decay as a potential therapeutic strategy in disorders caused by tRNA hypomodification.

## Introduction

Post-transcriptional epigenetic modifications of RNA—collectively known as the epitranscriptome—are increasingly recognized to exert diverse, widespread effects on gene regulation. More than 170 distinct RNA modifications have been cataloged to date, with approximately 80% occurring in tRNAs.^1,2^ tRNAs are transcribed as precursors by RNA polymerase III^3^ and then undergo a sequential maturation program that includes removal of 5′ leaders and 3′ trailers, splicing of intron-containing pre-tRNAs, post-transcriptional addition of the 3′-CCA terminus,^3,4^ and installation of numerous base and ribose modifications by dedicated tRNA-modifying enzymes.^5–7^ Upon completion of these processing steps, mature tRNAs are aminoacylated by their cognate aminoacyl-tRNA synthetases^8,9^ and delivered to the ribosome to decode mRNA codons during translation.^10^ Chemical modifications affect tRNA structure, dynamics, and interactions with other molecules to regulate their processing and stability, and ultimately translation.^5–7^ Many of these modifications are associated with human diseases.^6,11,12^ N^7^-methylguanosine (m^7^G) is a tRNA modification found in three domains of life.^13,14^ Methyltransferase-like 1 (METTL1) catalyzes the addition of m^7^G_46_ and interacts with the non-catalytic subunit WD repeat domain 4 (WDR4); both are widely conserved from yeast to mammals.^14–19^ The *Saccharomyces cerevisiae* mutants, *trm8Δ* and *trm82Δ*, both show loss of m^7^G_46_ modification and temperature sensitive growth.^13,20^ Growth defects at higher temperatures are more evident when other tRNA body modifications are lost.^21^ For example, *trm8Δ trm4Δ* double mutant have enhanced temperature sensitivity and a substantial reduction in the steady state level of tRNA(ValAAC).^21–23^ Moreover, synthetic approaches have identified factors involved in tRNA reduction and have led to the identification of a tRNA quality control system named the rapid tRNA decay (RTD) pathway, which degrades a subset of the tRNA species lacking any of several body modifications.^22–25^ Degradation of tRNAs by the RTD pathway is catalyzed by the 5′→3′ exonucleases, Rat1 and Xrn1, and is inhibited by a *met22Δ* mutation, which causes accumulation of the Met22 substrate, adenosine 3′, 5′ bisphosphate (pAp).^23,25–29^ The RTD is also observed in the fission yeast, *Schizosaccharomyces pombe*. *S. pombe trm8Δ* mutants show a temperature sensitive growth defect, which is primarily caused by decay of tRNA(TyrGUA) and to some extent tRNA(ProAAG) by the Rat1 ortholog, Dhp1. This indicates conservation of the overall RTD pathway, although there are differences in the tRNAs affected between the two species.^30,31^ In the *S. cerevisiae trm8Δ trm4Δ* mutant at 28°C and in the *S. pombe trm8Δ* mutant at 30°C, the amounts of tRNAs were reduced by about 30% compared with wild-type levels, indicating that the steady-state levels of hypomodified tRNAs decrease in physiological conditions.^21,31^ Indeed, tRNA(ValAAC) levels were recovered in *trm8Δ trm4Δ xrn1Δ* compared with those in *trm8Δ trm4Δ*, suggesting a potential involvement of the RTD pathway under physiological conditions. However, which tRNA metabolic steps contribute to the decrease in steady-state levels of unmodified tRNAs has not yet been investigated.^21,23^ A clear difference is observed for levels of aminoacylation of tRNAs between physiological and heat stress conditions. The majority of m^7^G-hypomodified tRNAs are deacylated under heat stress conditions, but not under physiological conditions.^21,22,25^ Therefore, even if the RTD pathway works under physiological conditions, it is unclear whether it is identical to that operating during heat stress conditions.

Another tRNA quality control system is the nuclear surveillance pathway that targets pre-tRNA(iMet) lacking m^1^A_58_.^32,33^ This pathway acts through the nuclear exosome TRAMP complex, which oligoadenylates the 3′ end of pre-tRNA(iMet), allowing its 3′–5′ exonucleolytic degradation by Rrp6 and Rrp44.^34–36^ While the nuclear surveillance pathway is well accepted, the RTD pathway has recently been suggested to play a more dominant role in maintaining pre-tRNA(iMet) in both *S. pombe* and *S. cerevisiae.*^34,37^ There is limited evidence for the tRNA quality control system acting on hypomodified tRNAs in animals. The RTD pathway plays important roles in tRNA(Leu) degradation under heat stress in mouse NIH3T3 cells.^38^ Moreover, in HeLa cells at 43°C, depletion of METTL1 and NSUN2 (a homolog of *S. cerevisiae* Trm4) reduced levels of tRNA(Val) in the presence of 5-fluorouracil, while tRNA(iMet) decay was mediated by XRN1 and XRN2.^39,40^ Although these reports suggest that the tRNA quality control system functions in mammalian cells under heat stress conditions, it remains unknown whether it also functions under physiological conditions.

Understanding the effect of tRNA quality control on hypomodified tRNA under physiological conditions is crucial because of its connection with animal development and human diseases.^6,11^ In *Drosophila mettl1* mutants at 25°C, a standard rearing temperature, the levels of tRNA(ValAAC) and tRNA(ProUGG) are significantly reduced in the testes, leading to ribosome collision and male sterility.^41^ This decrease in tRNA levels is much lower in other tissues,^41^ indicating that the effect of RNA modifications on tRNA abundance can vary among tissues. This possibility is supported by the involvement of tRNA modifications in human diseases. *METTL1* and *WDR4* are expressed in most tissues and developmental stages and their variants are involved in microcephalic primordial dwarfism, Galloway-Mowat syndrome,^42–44^ multiple sclerosis,^45^ male infertility,^46^ and tumorigenesis in acute myeloid leukemi,^47,48^ indicating a link between internal m^7^G regulation and pathological characteristics.^49^ Beyond m^7^G, deletion of FTSJ1, a 2′ -O-methyltransferase, lowers tRNA(Phe) levels and causes codon-specific translational deficits associated with behavioral abnormalities.^50,51^ In addition, NAT10–THUMPD1 (ac^4^C_12_) complex, TRMT1 (m^2,2^G_26_), TRMT10A (m^1^G_9_) and NSUN2 (m^5^C) have emerged as regulators of tRNA abundance in physiological conditions with potential links to human neurological disorders.^52–56^ The various physiological abnormalities of these disorders are observed in specific organs and cell-types despite the modifications occurring in all tissues. However, whether this arises from enhanced tRNA degradation is unclear.

We aimed to determine if tRNA quality control regulates the abundance of hypomodified tRNAs in higher eukaryotes under physiological conditions. We focused on m^7^G tRNA modification, which is associated with reduced levels of hypomodified tRNAs in *Drosophila*, mouse tissues, and human cell lines. First, we examined factors affecting hypomodified tRNA abundance in HCT116 cells, a human colorectal cancer cell line. We identified involvement of XRN2, whose depletion restores m^7^G hypomodified tRNA levels reduced by METTL1 knockdown; however neither XRN1 nor EXOSC10 (a yeast Rrp6 ortholog) were involved. To assess tRNA decay kinetics, we employed conditional degron systems to rapidly and reversibly deplete target proteins with two different inducers, 5-Ph-IAA and AGB1.^57,58^ Using this system, we found that hypomodified tRNA was reduced to a certain level without further reduction. This indicated the existence of two different steady states depending on the modification state of the tRNA. One is a state in which tRNA with m^7^G modification is present, and the other is a state in which only m^7^G-hypomodified tRNA is present. Moreover, re-expression of XRN2 after ligand removal promoted degradation of m^7^G-unmodified, indicating an XRN2-dependent RTD process under physiological conditions. Finally, we extended these findings *in vivo*: partial reduction of Rat1 (the *Drosophila* XRN2 ortholog) rescued Mettl1-dependent male sterility, indicating a conserved, physiologically meaningful genetic interaction between the m^7^G writer and the 5′→3′ exonuclease. Together, our results define a mammalian RTD pathway that actively eliminates m^7^G-hypomodified tRNAs under physiological conditions. From these findings, we suggest that modulation of tRNA decay has the potential to ameliorate diseases caused by tRNA hypomodification.

## Results

### Depletion of XRN2 restores the steady-state level of m⁷G-hypomodified tRNAs in METTL1 knockout HCT116 cells

To explore the underlying molecular mechanisms that control m⁷G-hypomodified tRNA levels, we anticipated that a system that can precisely and quickly control protein levels would be necessary. For this we used HCT116 cells, a human cell line derived from colorectal carcinoma. This cell line has recently been used in combination with the AID2 degron tag system that enables rapid and tunable control of target protein levels using small molecules to study the fundamental roles of genes that are essential to cell survival and replication processes.^57,59,60^ Depletion of METTL1 reduces steady-state levels of tRNAs in various human tumor cell lines(A549, LNZ308, HuCCT1, RBE, Huh7 and MHCC97H), but its effect in HCT116 has not been established.^48,61–63^ We therefore first analyzed m⁷G-modified tRNAs and tested whether METTL1 affects tRNA abundance in HCT116 cells. Using Cas9 and a guide RNA targeting exon 4 of METTL1 (Figure S1), we generated two different METTL1 knockout (KO) cell lines (METTL1KO1 and METTL1KO2). Western blot analyses confirmed the loss of ∼34 kDa METTL1 in METTL1KO1 and METTL1KO2 (Figure 1A). Levels of WDR4, a non-catalytic partner protein for METTL1, were only slightly decreased in METTL1KO cells, indicating that the stability of WDR4 was not significantly affected in the absence of METTL1 (Figure 1A). We then performed deep sequencing using tRNA reduction and cleavage sequencing (TRAC-seq)^19,64^ to investigate m^7^G-modified tRNAs in HCT116 cells. TRAC-seq identified 22 m^7^G-modified tRNAs in naive HCT116 cells (Figure S1B). The m^7^G-modified tRNAs had the RAGGU motif located in the variable loop (Figure S1C and S1D), as previously shown.^17,19,48,61^ Northern blotting showed that METTL1KO decreased the steady-state levels of tRNA(ValAAC) and tRNA(ProCGG/AGG/UGG), but not of tRNA(TrpCCA) (Figure 1B). Consistent with the northern blot data, RNA-seq analysis validated the reduced levels of tRNA(ValAAC) and allowed us to divide m^7^G-modified tRNAs into three groups according to steady-state levels of tRNA in METTL1KO cells (Figure 1C, 1D, S1E and S1F): destabilized in METTL1KO (Group 1), unaltered in METTL1KO (Group 2), and non-m^7^G-modified tRNAs (Group 3) (Figure 1B, 1D and Figure S1F), which is largely consistent with results from previous studies in mammalian cells and *Drosophila* testis.^41,61,62,65^ Together, these data indicate that METTL1 is required for m^7^G modification of a subset of tRNAs in HCT116 cells.

**Figure 1.**
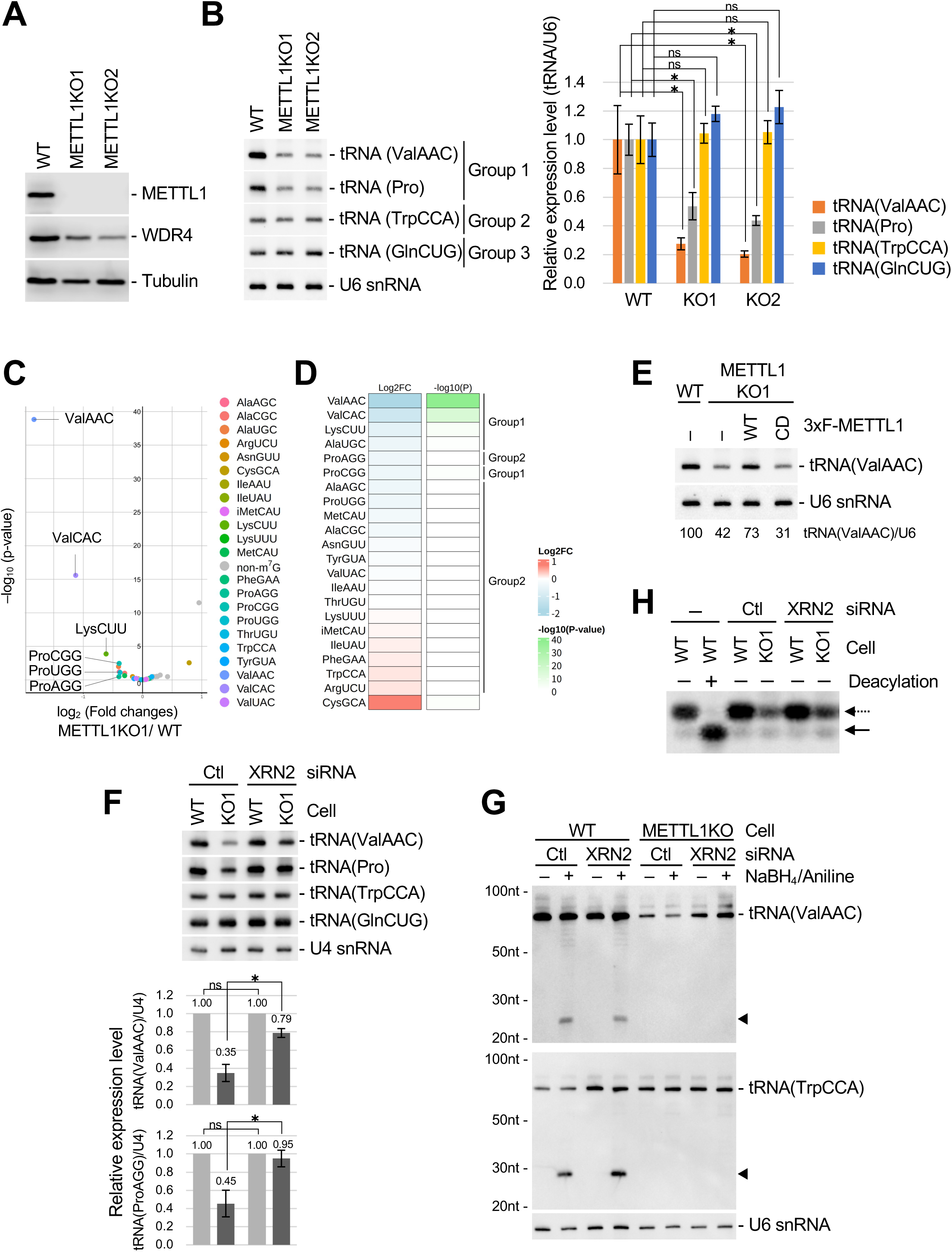
XRN2 knockdown suppresses METTL1 knockout-induced tRNA reduction. XRN2 depletion restores the abundance of m⁷G-hypomodified tRNAs in METTL1-deficient HCT116 cells. (A) Western blot analysis shows loss of METTL1 and a mild decrease in WDR4 levels in METTL1 knockout (KO) cells. (B) Northern blot (left) and quantification (right) showing reduced levels of tRNA(ValAAC) and tRNA(Pro) in METTL1KO1 and METTL1KO2 compared with wild-type (WT) HCT116 cells. U6 snRNA serves as a loading control. Bars represent mean ± s.d. from three independent experiments. Statistical significance was determined using an unpaired two-tailed Welch’s t-test (\**p* < 0.05; ns, not significant). The probe for tRNA(Pro) recognizes multiple isoacceptors, including tRNA(ProAGG), tRNA(ProCGG), and tRNA(ProUGG). (C) Volcano plot illustrating differential tRNA abundance between METTL1KO1 and WT cells. Each dot represents a single tRNA species; p-values were calculated using edgeR with Benjamini-Hochberg correction. (D) Heatmap showing changes in m^7^G-modified tRNA abundance between METTL1-KO (METTL1KO1) and WT HCT116 cells. tRNAs were classified into three Groups according to the difference in abundance between METTL1KO1 and WT, and p-values indicate significant differences; Group 1 (m^7^G-modified tRNAs and significantly decreased abundance, log_2_FC < 0, *p*< 0.05); Group 2 (other m^7^G-modified tRNAs, *p*≥ 0.05); Group 3 (non-m^7^G tRNAs). The *p*-values indicated correspond to those of (C). (E) Northern blot showing that re-expression of 3×FLAG-METTL1-WT, but not the catalytic-dead mutant (CD), restores tRNA(ValAAC) levels in METTL1-KO1 cells. Quantification revealed significant differences among the four groups (F(3,8) = 22.5, p = 0.0003, one-way ANOVA). Tukey’s post-hoc test showed that METTL1-WT significantly rescued tRNA levels compared with KO (*p* = 0.0407) and showed no significant difference from WT (*p* = 0.1036), whereas METTL1-CD failed to rescue. U6 snRNA serves as a loading control. See also Figures S1G and S1H for protein expression and quantification. (F) Northern blot showing that siRNA-mediated knockdown of XRN2 restores tRNA levels in METTL1KO1 cells, whereas control siRNA (Ctl) has no effect. The lower graphs show the relative expression levels of tRNA(ValAAC) and tRNA(Pro), quantified from the northern blot signals using U4 snRNA as a loading control. Data are representative of three independent experiments. Statistical significance was determined using an unpaired two-tailed Welch’s t-test (\**p* < 0.05, ns = not significant). The probe for tRNA(Pro) recognizes multiple isoacceptors, including tRNA(ProAGG), tRNA(ProCGG), and tRNA(ProUGG). (G) Northern blot of chemically treated total RNAs showing that the restored tRNAs remain m⁷G-unmodified. Probes hybridize to the 3′ ends of tRNA(ValAAC) and tRNA(TrpCCA); arrowheads mark cleaved fragments. (H) Acid-urea PAGE separating charged and uncharged tRNA(ValAAC) species from WT and METTL1KO1 cells. Deacylated RNA from WT cells (+ Deacylation) serves as a negative control. Dashed arrows represent aminoacylated tRNA(ValAAC), whereas solid arrows represent deacylated tRNA(ValAAC).

Transient re-expression of 3×FLAG-METTL1-WT, but not the catalytic-dead mutant (CD), in METTL1-KO1 cells restored tRNA(ValAAC) levels to near wild-type levels (Figure 1E, S1G and S1H), which is consistent with previous reports^19,41^. These results demonstrate that the catalytic activity of METTL1 is essential for maintaining tRNA abundance in HCT116 cells.

To gain insight into the potential involvement of tRNA degradation enzymes in the reduction of m^7^G-hypomodified tRNA levels, we tested whether components of the RNA exosome are involved in determining steady state levels of tRNAs. Strikingly, we found that depletion of XRN2 by siRNA-mediated knockdown (siKD) in METTL1KO cells caused the levels of Group 1 tRNAs(ValAAC and ProCGG) but not of Group 2 tRNA(TrpCCA) or Group 3 tRNA(GlnCUG) to recover (Figure 1F). In contrast, siKD of XRN1 or EXOSC10 had no effect on the recovery, indicating the specificity of XRN2 in regulating Group 1 tRNAs(ValAAC and ProCGG) in METTL1KO cells (Figure 1F and Figure S1J–O). To assess whether XRN2-depletion affects m^7^G-modification of tRNAs, we performed NaBH_4_/aniline treatment. As expected, depletion of XRN2 had no effect on the levels of m^7^G-modified tRNAs(ValAAC and TrpCCA), indicating that XRN2 did not affect METTL1 activity in HCT116 cells (Figure 1G). Critically, the recovered levels of tRNA(ValAAC) still lost m^7^G-modification, indicating that m^7^G-hypomodified tRNAs accumulate in METTL1KO/XRN2siKD cells.

We also examined aminoacylation levels of tRNAs in METTL1KO1 and METTL1KO1/XRN2KD cells by acid urea polyacrylamide gel electrophoresis (PAGE) and northern blot analysis. We found that the majority of tRNA(ValAAC) was aminoacylated in METTL1KO1 cells, which is consistent with previous data observed in *Drosophila mettl1* mutants.^41^ Moreover, tRNA(ValAAC) that are restored after XRN2 depletion are aminoacylated but not m^7^G-modified (Figure 1H), indicating that XRN2-depletion may rescue a functional tRNA in METTL1-depleted cells. Indeed, the proliferation rate of METTL1KO1 cells was comparable but slightly reduced compared with parental cells, indicating that the tRNA reduction was compatible with HCT116 cell proliferation (Figure S1I). This is consistent with the successful generation and proliferation of METTL1KO cell lines, despite the low levels of tRNA(ValAAC) and tRNA(ProCGG).

### Rapid degradation of METTL1 using the double degron system decreases tRNA levels

Depletion of XRN2 restored tRNA levels in METTL1KO cells; therefore, XRN2-mediated degradation of m^7^G-hypomodified tRNAs appears to be functional in HCT116 cells under physiological conditions. However, it is difficult to perform time-course analyses using METTL1KO or siRNA-based knockdown methods. To overcome this issue, we employed a rapid protein knockdown system that enables the direct degradation of target proteins induced by proteolysis-targeting short tags, named degron tags.^57,59,60^ We established a cell line in which a double-degron tag (mAB: <u>mA</u>ID-<u>B</u>romoTag)^57^ was inserted at the N-terminus of the endogenous METTL1 protein (mAB-METTL1) to enable rapid protein knockdown (pKD) (Figure 2A). Western blotting confirmed the rapid loss of mAB-METTL1 in the presence of 5-Ph-IAA and AGB1, with mAB-METTL1 undetectable within 30 min (Figure 2B). To check the status of m⁷G-modification, we performed NaBH₄/aniline treatment and found that the m⁷G-dependent cleavage fragment gradually reduced to ∼20% in the 48 h from mAB-METTL1 pKD (t_1/2_ = 26.8 h), indicating persistence of the modified tRNA for a long period of time (Figure 2C). We then checked tRNA abundance by northern blotting. The steady-state levels of tRNA(ValAAC) were reduced to ∼60% at 24 h, then further reduced to roughly ∼20% at 48 h, supporting that METTL1 depletion leads to a gradual loss of m⁷G modification accompanied by decreased tRNA abundance. (Figure 2C and 2D). Notably, the levels of tRNA at 48 h after mAB-METTL1 pKD are similar to those in METTL1KO1 and METTL1KO2 cells (Figure 1B) and no further decrease in tRNA levels was observed after 48 h, indicating that a decrease in tRNA levels initially occurs soon after mAB-METTL1 pKD and that the decrease may stop once a certain level is reached (Figure 2D). We next examined the aminoacylation status of tRNA(ValAAC). No accumulation of deacylated tRNA(ValAAC) was detected even after prolonged METTL1 depletion (Figure 2E). These results indicate that loss of m⁷G modification does not trigger tRNA deacylation.

**Figure 2.**
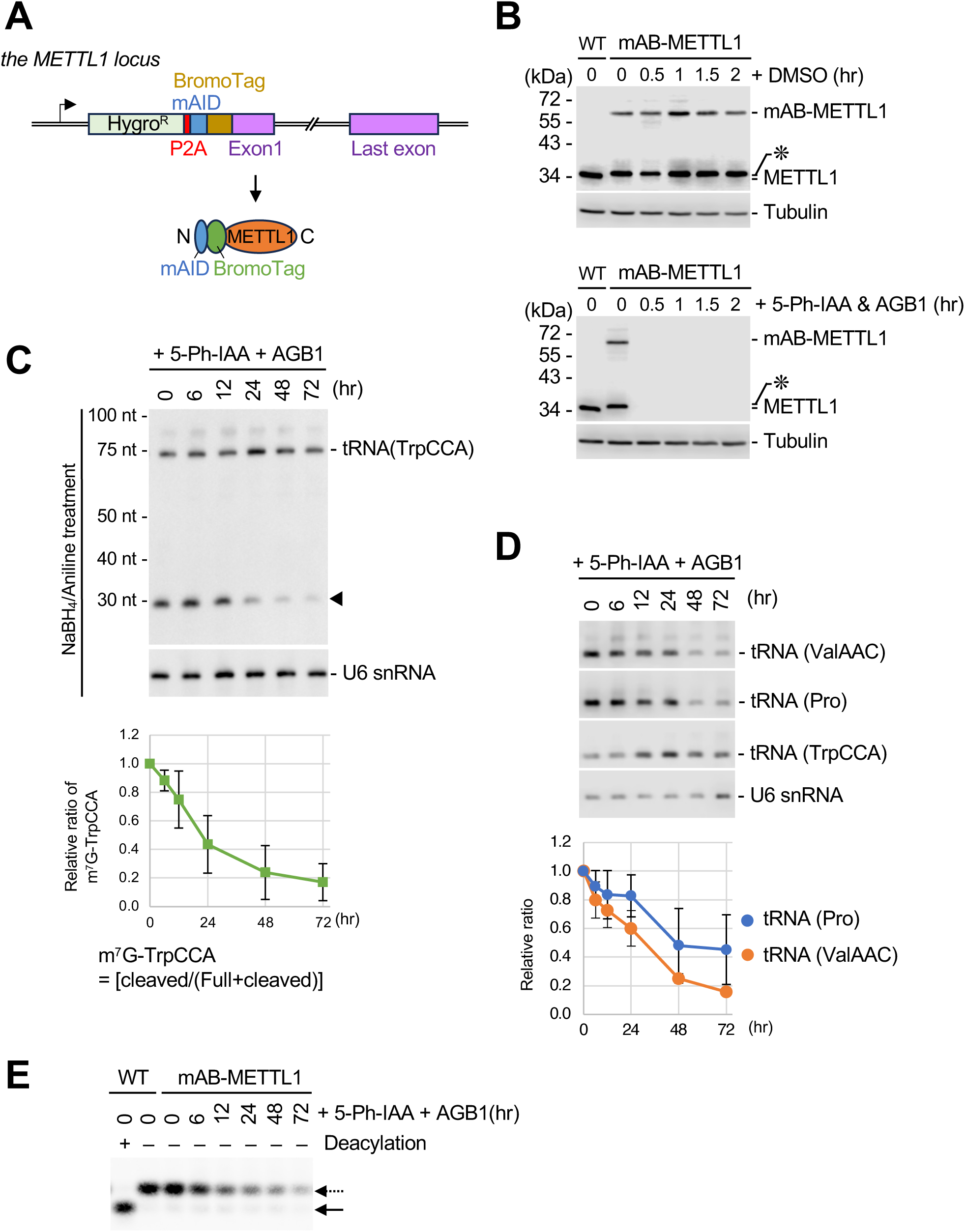
METTL1 depletion by the mAID-BromoTag system disrupts tRNA modification and stability. Acute METTL1 degradation via a dual-degron system reveals rapid tRNA decay kinetics. (A) Schematic illustration of mAID–BromoTag (mAB) insertion into the endogenous *METTL1* locus. (B) Western blot showing ligand-dependent degradation of mAB–METTL1 in HCT116 OsTIR1(F74G) cells treated with 1 µM 5-Ph-IAA and 0.5 µM AGB1. The asterisk indicates an additional band unique to mAB–METTL1 cells. (C) Northern blot analysis of NaBH_4_/aniline-treated total RNA showing time-dependent loss of m^7^G-modified tRNA(TrpCCA). Graph indicates relative m^7^G modification, normalized to time 0. (D) Northern blot showing the progressive reduction in total tRNA(ValAAC) and tRNA(Pro) abundance following METTL1 depletion. The graph quantifies relative tRNA levels normalized to time 0; U6 snRNA serves as a loading control. Data are representative of three independent experiments. The probe for tRNA(Pro) recognizes multiple isoacceptors, including tRNA(ProAGG), tRNA(ProCGG), and tRNA(ProUGG). (E) Aminoacylation analysis of tRNA(ValAAC) using the same RNA samples as in (C) and (D). Deacylated RNA from WT cells (+ Deacylation) serves as a negative control. Dashed arrows represent aminoacylated tRNA(ValAAC), whereas solid arrows represent deacylated tRNA(ValAAC).

### XRN2-dependent degradation of m⁷G-hypomodified tRNAs

Our observations of XRN2siKD restoring steady state levels of Group 1 tRNAs in METTL1KO cells (Figure 1F) and of mAB-METTL1 loss immediately reducing tRNA(ValAAC) (Figure 2D) indicate that m⁷G-hypomodified tRNAs are susceptible to degradation by a mechanism similar to the rapid tRNA decay (RTD) pathway. However, METTL1 depletion by the degron system (mAB-METTL1pKD) gave rise to mixed pools of modified and unmodified tRNAs (Figure 2C), making it difficult to accurately measure the levels of unmodified tRNAs alone. To overcome this issue, we tagged both METTL1 and XRN2 with distinct degron tags, BromoTag and mAID, respectively (Figure 3A). Following AGB1 addition (for METTL1pKD), we observed the expected reduction in tRNA levels (ValAAC and ProCGG/AGG/UGG) (Figure 3C, lane 3). Addition of both AGB1 and 5-Ph-IAA (for METTL1-pKD and XRN2-pKD) restored tRNA levels (Figure 3C; compare lane 3 vs lane 4), which is consistent with the knockdown experiment (Figure 1F) where the restored tRNAs were m⁷G-unmodified (Figure 1G). To test whether the recovered tRNAs are susceptible to degradation by XRN2, we washed out 5-Ph-IAA from the medium (time 0) to restore XRN2 levels. After wash out of 5-Ph-IAA and maintenance of AGB1 (time 0 in Figure 3B), XRN2 increased within ∼12 h (Figure 3C; lanes 5–8, 10–13) to levels comparable to those in wild-type cells. In METTL1pKD cells, tRNA levels (ValAAC and ProCGG/AGG/UGG) decreased after XRN2 recovery with t_1/2_ ≈ 24 h (lanes 4–8). However, no decrease was observed upon XRN2 recovery in the presence of METTL1 (lanes 10–13), indicating that the tRNA reduction depends on the m^7^G-modification status. All kinetics were measured in the presence of an RNA polymerase III inhibitor, confirming transcription-independent reduction of m⁷G-modified Group 1 tRNAs(ValAAC and ProCGG) but not of Group 3 tRNA(GlnCUG). Considering these observations together, we propose that XRN2 is responsible for the degradation of m⁷G-hypomodified tRNAs in HCT116 cells under physiological conditions (Figure 3C).

**Figure 3.**
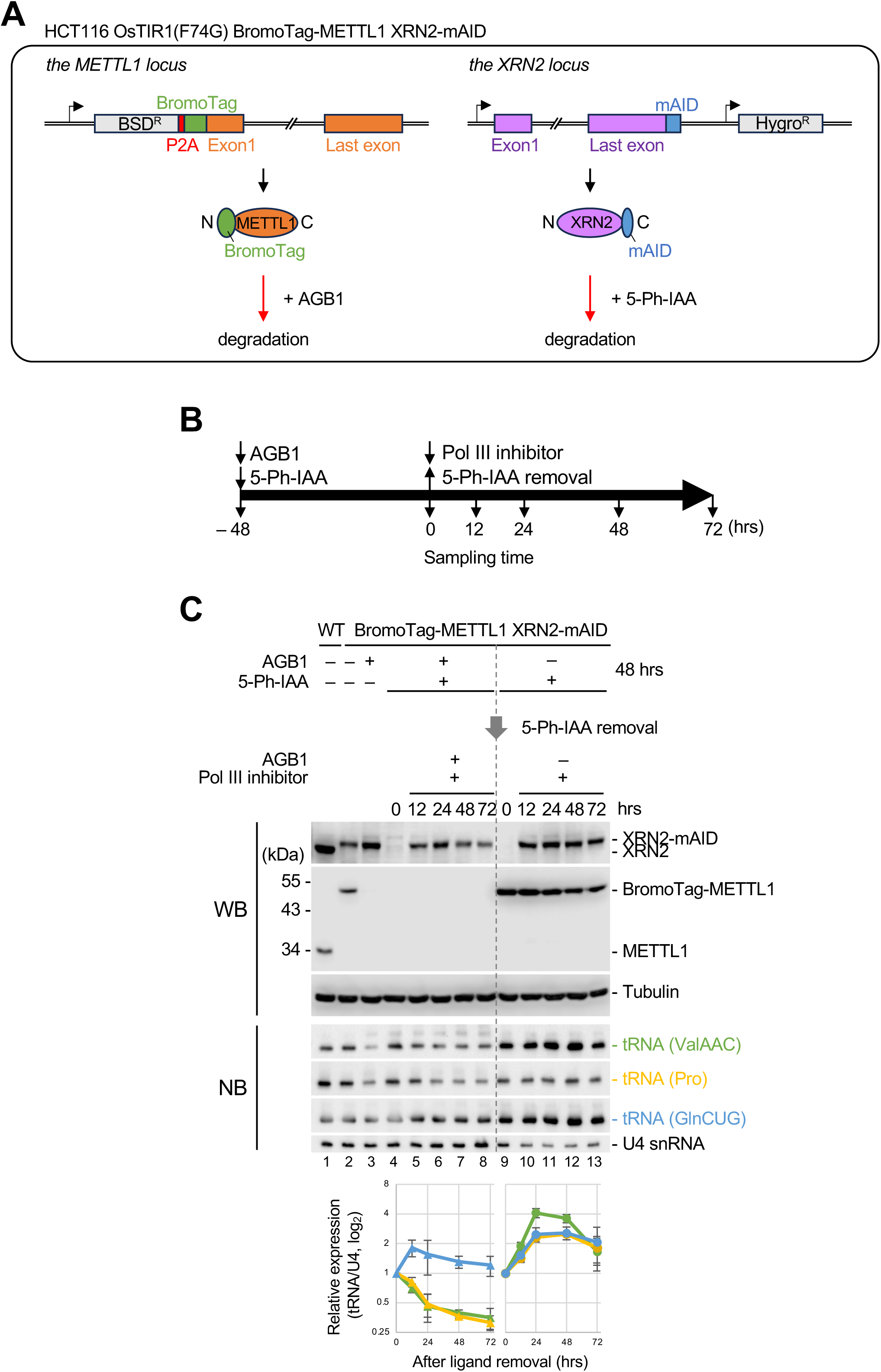
XRN2 recovery reveals its role in regulating m⁷G-hypomodified tRNA degradation. Re-expression of XRN2 selectively eliminates m⁷G-hypomodified tRNAs under physiological conditions. (A) Schematic diagram of the generation of a cell line harboring BromoTag-METTL1 and XRN2-mAID. (B) Experimental workflow illustrating ligand treatments and sampling time points for METTL1 depletion and XRN2 recovery. (C) Western and northern blot analyses showing that re-expression of XRN2 triggers the decay of hypomodified tRNAs. Top, western blot (WB) for METTL1 and XRN2. Middle, northern blot (NB) for tRNA(ValAAC) and tRNA(Pro). Bottom, quantification of relative tRNA levels from three independent experiments. U4 snRNA serves as a loading control. The probe for tRNA(Pro) recognizes multiple isoacceptors, including tRNA(ProAGG), tRNA(ProCGG), and tRNA(ProUGG).

### Rat1 (an XRN2 homolog) genetically interacts with Mettl1 in *Drosophila*

Finally, to test whether the relationship between METTL1 and XRN2 for tRNA degradation is genetically relevant, we focused on the critical role of *Drosophila* Mettl1 in male fertility.^41^ Based on sequence similarity, *Drosophila* Rat1 is predicted to be a homolog of budding yeast Rat1 and metazoan XRN2. A *rat1^SK3^*mutant harboring an N-terminal single-base deletion is predicted to eliminate functional Rat1 (Figure S2). *rat1^SK3^* is homozygous lethal, consistent with essential roles of XRN2/Rat1 in eukaryotes (data not shown). To assess a genetic interaction between Mettl1 and Rat1, we utilized a fly heterozygous for a loss of function *rat1* mutation (*rat1^SK3^*/+), since *Drosophila* has been used as a powerful tool to investigate genetic interactions of essential genes with great sensitivity, for example, using heterozygotes to discover chromatin modifiers.^66^ We generated *mettl1^KO1^; rat1^SK3^/+* males and crossed them to wild-type females. *mettl1^KO1^* males produced ∼0 progeny; however, *mettl1^KO1^; rat1^SK3^/+* males restored progeny counts, comparable to those of *yw* and *rat1^SK3^*/+ controls (Figure 4A), indicating genetic interaction between *mettl1* and *rat1*. A similar genetic suppression was observed with *wuho*^67^, the *Drosophila* ortholog of human WDR4 (Figure 4B), indicating that developmental defects caused by loss of m^7^G modification machinery are alleviated by downregulation of *rat1*.

**Figure 4.**
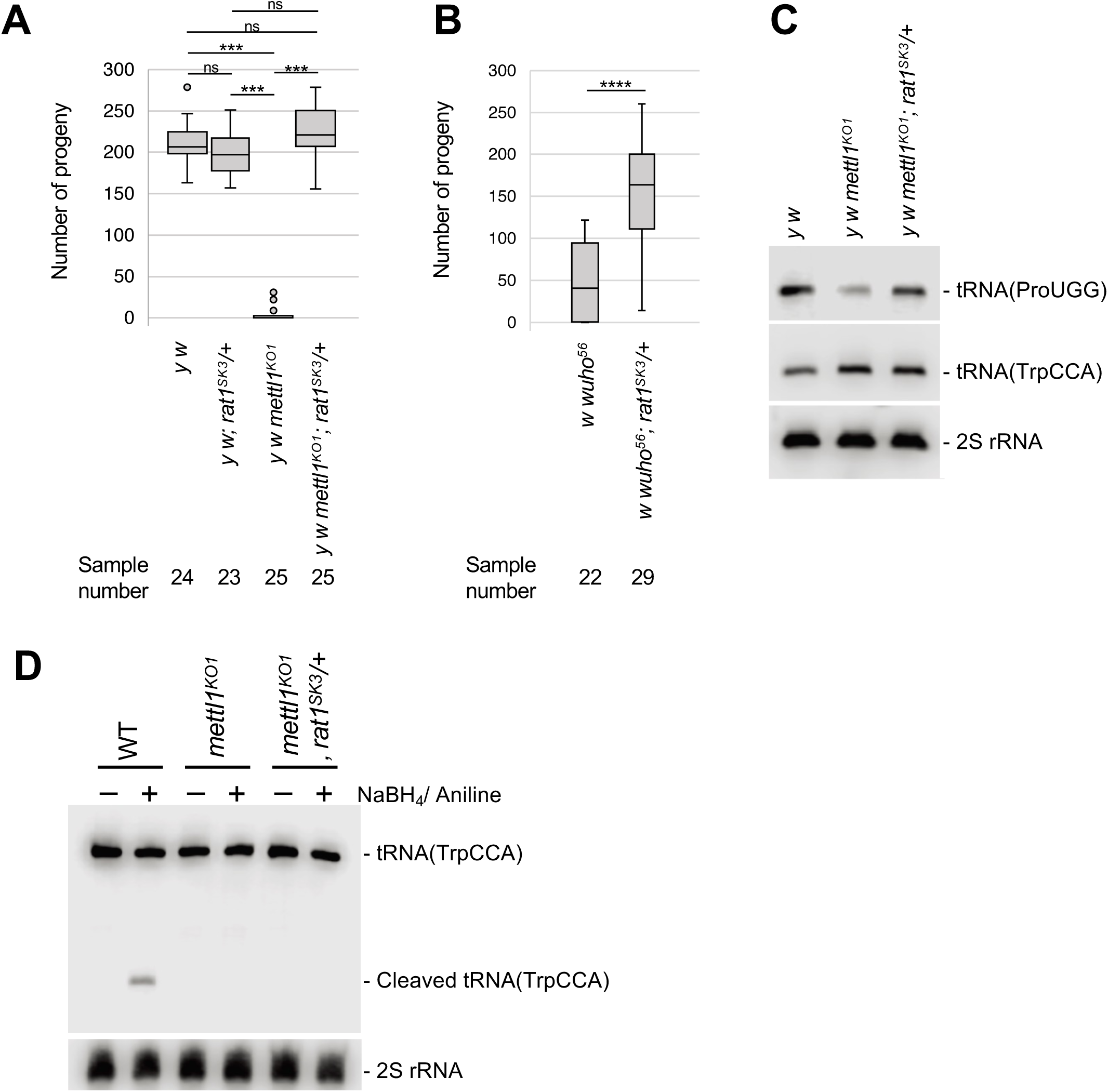
Rat1/XRN2 genetically interacts with Mettl1 in *Drosophila*. Partial reduction of Rat1 restores fertility and tRNA abundance in *mettl1 mutant* flies. (A) Fertility assay showing progeny numbers per male for *y w*, *y w; rat1 /+*, *y w mettl1^KO1^*, and *y w mettl1^KO1^; rat1^SK3^/+*. *y w mettl1^KO1^* males showed almost complete sterility, whereas *y w mettl1^KO1^; rat1^SK3^/+* males displayed a significant rescue to near wild-type levels. One-way ANOVA revealed a significant difference among genotypes (F(3, 92)=279.5, *p*<0.0001); Tukey’s HSD test showed \*\*\**p*<0.001 for both *mettl1^KO1^*vs *mettl1^KO1^; rat1^SK3^/+* and *mettl1^KO1^*vs *y w* comparisons. Boxes represent the interquartile range; whiskers show minimum–maximum; dots represent biological replicates (mean ± s.d.). (B) Similar fertility rescue in *wuho^56^; rat1^SK3/+^* males. Boxplot showing fertility of *w wuho^56^* and *w wuho^56^; rat1^SK3^*/+ males. Partial reduction of rat1 significantly restored fertility in *wuho^56^*mutants (t = -6.85, \*\*\*\**p* = 1.1 × 10⁻⁸; unpaired two-tailed Welch’s t-test). Bars represent individual data points (n = 22 for *w wuho^56^*, n = 29 for *w wuho^56^; rat1^SK3^*/+). (C) Northern blot analysis comparing tRNA levels in testes from WT (*yw*), *mettl1* knockout (*mettl1^KO1^*), and *mettl1^KO1^* with a heterozygous *rat1* mutation (*mettl1^KO1^; rat1^SK3^/+*). tRNA(ProUGG) levels were increased in *mettl1^KO1^; rat1^SK3^/+* relative to *mettl1^KO1^*, indicating that partial reduction of Rat1 restores tRNA abundance in *mettl1* mutant. In contrast, tRNA(TrpCCA), which is also m⁷G-modified by METTL1 but shows no abundance change upon METTL1 loss, remained unaffected. 2S rRNA serves as a loading control. (D) Northern blot of NaBH₄/aniline-treated RNA from testes showing that restored tRNAs remain m⁷G-unmodified. Probes hybridize to the 3′ end of tRNA(TrpCCA). Boxplots show median, quartiles, and range; numbers below indicate sample size.

To confirm if the genetic interaction between Mettl1 and Rat1 is associated with tRNA levels, we examined tRNA abundance in testes and found increased levels of tRNA(ProUGG) in *mettl1^KO1^*; *rat1^SK3^/*+ relative to those in *mettl1^KO1^* (Figure 4C). Notably, NaBH₄/aniline treatment showed no restoration of m⁷G in *mettl1^KO1^*; *rat1^SK3^/+* males, indicating that m⁷G-unmodified tRNA levels had increased, which rescued fertility. Partial reduction of Rat1/XRN2 therefore mitigates the organismal consequences of Mettl1 loss, supporting the concept that inhibiting the RTD pathway, in addition to restoring modifications, may be a tractable strategy for diseases caused by tRNA hypomodification.

## Discussion

Our study identifies XRN2 as the central nuclease responsible for degrading m⁷G-hypomodified tRNAs in mammalian cells under normal physiological conditions. Loss of METTL1, either by knockout or degron-mediated acute depletion, rapidly reduced the abundance of m⁷G-modified tRNAs, establishing a robust model for studying tRNA decay (Figures 1A–D, S1E, S1F, and 2A–D). RNAi experiments indicated that XRN2 is specifically involved in the decrease of m⁷G-hypomodified tRNA abundance, whereas XRN1 and EXOSC10 are not (Figures 1F, 1G, and S1J–O). Using degron-based acute protein depletion combined with time-resolved decay analysis, we demonstrated that re-expression of XRN2 selectively degrades m⁷G-hypomodified tRNAs, providing direct evidence that XRN2 discriminates tRNAs based on their modification status (Figure 3A–C). Furthermore, we found that reduction of Rat1/XRN2 rescued *Drosophila* male sterility caused by Mettl1 loss (Figure 4A–D). Together, our degron-based and genetic approaches reveal that the animal RTD pathway functions under physiological temperatures and is important for tRNA reduction even outside of heat stress conditions. This discovery is expected to contribute to elucidating the pathogenesis of diseases caused by the loss of not only m^7^G but also various other tRNA modifications, and to developing therapeutic approaches to overcome RNA modopathies, the class of diseases and disorders caused by aberrations in tRNA modifications.

The RTD pathway was first characterized in yeast, where Rat1/Xrn2 and Xrn1 cooperatively degrade hypomodified tRNAs.^21,23,31^ Moreover, the previous findings of the RTD pathway in HeLa and NIH3T3 extends this concept to mammals.^38,39,40^ While, those studies probed accelerated decay after heat shock, however, it remains unclear whether RTD functions constitutively under physiological conditions. This is an important step toward understanding the pathogenesis of various diseases caused by alterations of tRNA modification status, because these diseases occur at normal temperatures but not under heat stress. Our study demonstrates that RTD functions constitutively under physiological conditions, independent of stress. Thus, RTD should therefore be viewed as a housekeeping pathway that continuously monitors tRNA integrity, rather than as a stress-induced response. Notably, in *S. cerevisiae* that *trm8Δ* t*rm4Δ* strain exhibit reduced tRNA abundance and *trm8Δ trm4Δ rat1-107 xrn1Δ* strain under permissive temperatures (28–30°C), even before heat stress.^23^ It seems likely that RTD operates at non-heat stress conditions in *S. cerevisiae*. Furthermore, because similar reduction of the steady-state levels of tRNAs has been observed for other hypomodified tRNAs, such as those lacking ac^4^C, m^2,2^G, m^1^G or m^5^C, RTD appears to represent a general surveillance network that recognizes multiple classes of defective tRNAs.^52–56^ These findings collectively support the existence of a modification–decay axis that selectively eliminates aberrant tRNAs from the translational pool.

We propose that RTD functions as part of a broader tRNA modification–decay homeostat, a dynamic system balancing tRNA modification, abundance, and translation efficiency. In this framework, tRNA stability is not a passive consequence of modification loss but a regulated outcome of modification-dependent surveillance. When modification enzymes, such as METTL1, are compromised, unmodified tRNAs are rapidly recognized and eliminated by RTD to prevent their aberrant incorporation into the translation machinery. This principle parallels mRNA and rRNA surveillance systems and indicates that tRNA metabolism is governed by a similar homeostatic circuit that preserves proteome quality.^6,7^ This model also accommodates cross-talk among tRNA modifications, in which loss of one modification can influence the stability or installation of others. For instance, the absence of m^7^G in the variable loop may perturb local structure and thereby affect nearby modifications, such as m^5^C or ac^4^C, cumulatively enhancing decay susceptibility.^13^ RTD therefore likely serves as a final checkpoint in the integration of multiple epitranscriptomic cues, converting complex modification patterns into binary decay decisions.

Mechanistically, our dual-degron analysis provides direct evidence that the RTD pathway can operate on aminoacylated tRNAs lacking proper modification. Acute depletion of METTL1 led to a time-dependent decrease in tRNA abundance without detectable accumulation of deacylated species, indicating that tRNA decay can proceed independently of prior deacylation (Figure 2E). This contrasts with the canonical yeast RTD model, in which deacylation exposes the 5′ end for exonucleolytic attack.^22,25^ In mammalian cells, the absence of m^7^G modification itself appears to serve as the primary decay trigger, allowing the surveillance machinery to recognize structural defects even in aminoacylated tRNAs. Consistent with this, overexpression of EF1α mitigated the reduction of m^7^G -deficient tRNAs upon METTL1 depletion^22,41,65^, suggesting that these unmodified yet aminoacylated tRNAs can transiently engage the translation machinery. We propose that RTD kinetically competes with translation: EF1α binding transiently shields tRNAs, whereas dissociation exposes unmodified species to XRN2-mediated decay. This dual control ensures that only functional tRNAs remain engaged in translation. Notably, a methyltransferase-independent role of METTL1 in promoting tRNA aminoacylation and translational control has been reported in other contexts.^68^ However, in our HCT116 system, METTL1 depletion did not measurably alter aminoacylation of the affected tRNAs (Figures 1H and 2E), indicating that such activity is not detectable in this cellular context and is dispensable for the XRN2-dependent decay we observed. These observations reinforce the idea that, in human cells, loss of m^7^G modification *per se* is sufficient to trigger RTD, independent of auxiliary effects on tRNA charging. Another notable feature of our data is that m⁷G-deficient tRNA(ValAAC) persists at about 20% of wild-type levels in METTL1pKD cells. This indicates a selectively protected subpopulation that escapes decay, probably because of stable aminoacylation and ribosomal engagement. Such dual gating—modification and aminoacylation—may allow cells to balance translational capacity against stringent quality control, ensuring proteome integrity even under partial modification defects.

The conservation of RTD from yeast to mammals illustrates a simple but powerful rule of RNA homeostasis; defective tRNAs are not tolerated, even under normal conditions. Nevertheless, the affected tRNA species differ among species, indicating that RTD has co-evolved with lineage-specific tRNA repertoires and translational demands. The partial rescue of Mettl1-dependent male sterility by heterozygous Rat1 reduction in *Drosophila* further indicates that modulating RTD activity can buffer developmental phenotypes^41^ and indicates that tRNA decay is a tunable process with physiological relevance.

From a biomedical perspective, our results raise the possibility that stabilizing hypomodified tRNAs could be as beneficial as restoring the modification itself. Excessive RTD activity may underlie pathological tRNA loss in disorders such as microcephaly, neurodevelopmental syndromes, and infertility linked to METTL1 or WDR4 mutations.^42,43^ Conversely, previous studies showed that dysregulated METTL1 activity promotes oncogenic transformation and suggested that both insufficient and excessive control of the METTL1–tRNA axis can disrupt cellular homeostasis.^48,61,62^ Targeting the decay machinery—rather than modification enzymes—may therefore offer a viable intervention strategy. Pharmacological or proteolysis targeting chimeric (PROTAC)-based inhibition of 5′→3′ exonucleases or RNA helicases may restore tRNA balance and translational homeostasis. Indeed, recent advances in PROTAC technology demonstrate that targeted protein degraders can achieve more sustained and selective suppression of enzyme activities than conventional small-molecule inhibitors,^69,70^ offering a compelling avenue for “anti-decay” therapeutics in tRNA modification disorders.

### Limitations and future perspectives

Our study demonstrates that XRN2 mediates the decay of m^7^G-hypomodified tRNAs under physiological conditions; however, several key questions remain. The precise molecular mechanism by which XRN2 distinguishes modified from unmodified tRNAs, and whether auxiliary factors such as helicases or adaptor proteins assist this process, remain to be elucidated. Notably, we showed that the reduction of XRN2 but not XRN1 suppressed the decrease in tRNA levels caused by METTL1 loss (Figures 1F, S1K, S1M, and S1N). This is stark contrast to recent finding in a heat-treated condition in NIH3T3, where loss of either XRN1 or XRN2 alone had no effect, but loss of both XRN1 and XRN2 recovers tRNA levels, suggesting that XRN2 and XRN1 have overlapping roles in a heat shock-induced RTD pathway.^38^ Thus, the factors involved differ between heat stress and physiological conditions, also suggesting that the molecular mechanisms underlying RTD may differ.

Our analyses were performed in human cultured cells and *Drosophila* testes. Previous work showed that loss of Mettl1 in *Drosophila* causes obvious abnormality restricted to the testis^41^, indicating tissue-specific vulnerability to tRNA hypomodification. It will therefore be crucial to determine whether similar RTD-dependent regulation operates in other mammalian tissues or cell types, particularly in those affected in human diseases associated with METTL1 or WDR4 mutations, such as neural, hepatic, and germline tissues. Investigating how RTD activity correlates with cell-type-specific translational demands may reveal why tRNA modification disorders manifest in distinct organs. Studies combining biochemical, single-cell, and clinical approaches are needed to define how RTD contributes to the tissue specificity and pathogenesis of tRNA-related diseases.

## Acknowledgments

We are grateful to T. Miyoshi, A. Masuda, Y. Kitahara and N. Chalkarova for technical assistance and the Bloomington Drosophila Stock Center and the Drosophila Genetic Resource Center in Kyoto Institute of Technology for providing the fly strains. We also thank Jeremy Allen, PhD, from Edanz (https://jp.edanz.com/ac) for editing a draft of this manuscript.

This work was supported by JSPS KAKENHI under grant numbers, JP25K09570 (KM), JP22H02669 (KS), JP23K23932 (KS) and JP25K02209 (KS) and the Takeda Science Foundation (KS).

## Data and code availability

The RNA-seq data of small RNAs have been deposited with links to BioProject accession number PRJDB37935 in the DDBJ BioProject database.

## EXPERIMENTAL MODEL AND STUDY PARTICIPANT DETAILS

### Cell lines

HCT116 cells (human colorectal carcinoma; ATCC, CCL-247) were cultured in McCoy’s 5A medium supplemented with 10% fetal bovine serum, 2 mM L-glutamine, 100 U/mL penicillin, and 100 μg/mL streptomycin at 37°C in a humidified incubator with 5% CO_2_. All HCT116 cell lines expressing degron-fused proteins were generated as previously described.^71^ Briefly, donor and CRISPR plasmids were constructed for tagging each gene of interest. Cells were co-transfected with a donor plasmid and a CRISPR plasmid and subsequently selected in the presence of the appropriate antibiotic. Single colonies were isolated, and clones containing bi-allelic insertions at the target gene loci were identified by genomic PCR. The production of degron-fused target proteins was confirmed by western blotting. To generate METTL1 knockout HCT116 cells, single-guide RNAs (sgRNAs) targeting *METTL1* were cloned into the pX330 vector.^72^ HCT116 cells were transfected with the sgRNA-expressing vector (sgRNA5) and a puromycin resistance plasmid using Viafect (Promega, #E4981). One day after transfection, cells were seeded at low density and selected with 1 μg/mL puromycin for 24 h. Single colonies were isolated approximately 1 week later and screened by PCR and/or western blotting using an anti-METTL1 antibody (ProteinTech, 14994-1-AP) to identify knockout clones.

### *Drosophila* strains

A complete list of fly stocks used in this study is provided in Supplementary Table 2. All stocks were maintained at 25°C on standard medium. *Mettl1* knockout (*Mettl1^KO1^*) lines were described previously.^41^ The *rat1* mutant line (*rat1^SK3^*) generated using the transgenic CRISPR-Cas9 method^73^ was obtained from the National Institute of Genetics Fly Stock Center (#M2L-0047, Mishima, Japan). Strain *w* wuho*^56^*/FM7c*, *sn*^+^ was obtained from Bloomington *Drosophila* Stock Center (#41112).

### Study participants

No human subjects were involved in this study.

## METHOD DETAILS

### Plasmid constructions

The pET30a-AlkB (Addgene #79050) and pET30a-AlkB-D135S (Addgene #79051) plasmids were described previously.^74^ The Hygro-P2A-mAB (pMK462; Addgene #79051) and BSD-P2A-BromoTag (pMK445; Addgene #214375) plasmids were previously described in Hatoyama et al.^57^ The pBS-mAID-Hygro (pMK287; Addgene #72825) plasmid was described previously.^59^ The CRISPR-Cas9 backbone plasmid is pX330-U6-Chimeric_BB-CBh-hSpCas9 (Addgene #42230).^72^

Plasmids generated in this study for the CRISPR/Cas9-mediated knockout or knock-in system include pX330-U6-Chimeric_BB-CBh-hSpCas9_hMETTL1_sgRNA1 (pKMn811), pX330-U6-Chimeric_BB-CBh-hSpCas9_hMETTL1_sgRNA5 (pKMn852), pX330-U6-Chimeric_BB-CBh-hSpCas9_hXRN2_sgRNA-C2 (pKMn930), pBS-Hygro-P2A-mAB-hMETTL1_Narm (pKMn805), pBS-BSD-P2A-BromoTag-hMETTL1_N (pKMn924), and pBS-hXRN2_Cterm_mAID-Hygro (pKMn932). These CRISPR plasmids expressing sgRNA were designed to target human METTL1 (sgRNA1: 5′- TCCGGCCACGTTCCGAGTCT/CGG-3′, sgRNA5: 5′- CGTAGGCATATTCTGCTAGC/AGG-3′), XRN2 (5′- TGTAGAATGATGAAAGGATT/TGG-3′).

Expression constructs generated in this study include pcDNA3.1-3×FLAG-hMETTL1-WT (pKMn854) and pcDNA3.1-3×FLAG-hMETTL1-Catalytic Dead (CD) (pKMn1033). The catalytic-dead (CD) mutation was designed based on Lin et al.^19^ All newly generated constructs were verified by Sanger sequencing.

### Antibodies

For immunoblotting, the following primary antibodies were used: anti-METTL1 polyclonal (ProteinTech, 14994-1-AP), anti-WDR4 rabbit polyclonal (Abcam, ab169526), anti-β-Tubulin mouse monoclonal (Developmental Studies Hybridoma Bank, E7), anti-XRN2 rabbit polyclonal (ProteinTech, 112671-1-AP), anti-XRN1 rabbit polyclonal (ProteinTech, 23108-1-AP), and anti-EXOSC10 rabbit polyclonal (ProteinTech, 11178-1-AP). HRP-conjugated anti-rabbit IgG (Cell Signaling, #7074) and HRP-conjugated anti-mouse IgG (Thermo Fisher Scientific, #32230) were used as secondary antibodies.

### Plasmid transfection

HCT116 cells were seeded at a density of 1.0×10^5^ cells per well in 6-well plates 24 h before transfection. Cells were transfected with 25 ng of plasmid DNA (pcDNA3.1-3×FLAG-hMETTL1-WT or pcDNA3.1-3×FLAG-hMETTL1-CD) using ViaFect Transfection Reagent (Promega, #E4981) according to the manufacturer’s protocol. After 96 h, total RNA and protein were extracted for northern and western blot analyses, respectively.

### siRNA transfection

HCT116 cells were transfected with siRNAs using Lipofectamine RNAiMAX (Thermo Fisher Scientific, #13778-075) according to the manufacturer’s instructions. Briefly, cells were seeded at a density of 2 × 10^5^ cells per well in a 6-well plate 24 h before transfection. Cells were transfected with siRNAs at a final concentration of 10 nM and plated in Opti-MEM (Thermo Fisher Scientific, #31985-070). After incubation for 48 h, cells were harvested for subsequent analyses. The siRNA sequences used are listed in Supplementary Table 1. MISSION siRNA Universal Negative Control #1(Sigma-Aldrich, SIC001) was used as a negative control.

### Degrader compounds

5-Ph-IAA was synthesized as previously described.^60^ AGB1 was commercially obtained (Tocris #7686).

### Inducible degradation of METTL1 and XRN2

To generate the degron cell lines (mAB-METTL1, BromoTag-METTL1, and XRN2-mAID), degron tags were introduced by CRISPR-mediated knock-in.

For inducible degradation of mAB-METTL1, OsTIR1(F74G)-expressing HCT116 cells were used, and degron activity was induced with 1 μM 5-Ph-IAA (mAID ligand) and 0.5 μM AGB1 (BromoTag ligand). For inducible degradation of BromoTag-METTL1, the BromoTag system was used with 0.5 μM AGB1. For inducible degradation of XRN2-mAID, the mAID system was used with 1 μM 5-Ph-IAA. Cells were treated with the ligands for the indicated times before downstream analyses. To re-express XRN2, 5-Ph-IAA was removed from the culture medium, and cells were collected at the indicated time points. To monitor the degradation of m⁷G-hypomodified tRNAs upon XRN2 re-expression, transcription by RNA polymerase III was inhibited by adding 30 μM ML60218 (MedChemExpress, #HY122122).

### Western blotting

Western blotting was performed as described previously^75^, with minor modifications.

### Northern blotting

Northern blotting was performed using a modified method based on previously described protocols.^41,76,77^ Total RNAs were isolated using ISOGEN (Nippon Gene, #319-90211) following the manufacturer’s instructions. For general detection of tRNAs, 2 µg of total RNA was denatured with Gel Loading Buffer II (Thermo Fisher Scientific, #AM8546G) for 3 min at 95°C and resolved by electrophoresis on 12% urea–PAGE gels containing 7 M urea, 10 × TBE (1 M Tris base, 1 M boric acid, 0.02 M EDTA), and 12% acrylamide. To assess m^7^G-dependent cleavage and to confirm tRNA abundance, 2 µg of total RNA was resolved by electrophoresis and transferred to Hybond N+ membranes (Cytiva). Transferred RNAs were UV crosslinked (1200 × 100 mJ/cm² at 254 nm) and prehybridized in hybridization buffer (7% SDS, 0.2 M sodium phosphate, pH 7.2, 1 mM EDTA). Membranes were hybridized with DIG-labeled DNA probes (Merck, #3353575910) in hybridization buffer at 42°C overnight, washed with 1 × SSC, blocked using the DIG Wash and Block Buffer Set (Merck, #11585762001), and incubated with anti-DIG alkaline phosphatase Fab fragments (Merck, #11093274910). Signals were detected using CDP-Star (Merck, #12041677001) and imaged with a ChemiDoc Touch (Bio-Rad). Band intensities were quantified using Image Lab software (Bio-Rad). The DNA probe sequences used are listed in Supplementary Table 2.

### m^7^G site-specific reduction and cleavage

m^7^G-dependent RNA cleavage was performed by incubating 2.5 µg of total RNA with 0.1 M sodium borohydride (NaBH₄) and 1 mM free m^7^GTP on ice for 30 min in the dark.

Reduced RNAs were precipitated with 3 M sodium acetate (pH 5.2, Thermo Fisher Scientific), glycogen (Nacalai Tesque), and ethanol at −20°C for at least 1 h. RNA samples reduced by NaBH₄ were subsequently incubated with an aniline-acetate solution (H₂O:glacial acetic acid:aniline, 7:3:1) at room temperature for 2 h in the dark to induce m^7^G site-specific cleavage, as described previously.^78,79^ The resulting RNA fragments were analyzed by northern blotting.

### TRAC-seq and profiling of tRNA expression

Sequencing and data analysis were performed as previously described^64^ with minor modifications. Small RNAs (<200 nt) were isolated from total RNA using a mirVana miRNA Isolation Kit (Thermo Fisher Scientific, AM1561) according to the manufacturer’s instructions. To remove methylation that inhibits reverse transcription of tRNAs, 2.5 µg of extracted small RNAs was treated with recombinant ALKB and ALKB D135S proteins. Demethylated small RNAs were subjected to reduction and cleavage as described in the “m^7^G site-specific reduction and cleavage” section. After cleavage and ethanol precipitation, small RNA sequencing libraries of m^7^G tRNAs were prepared using a NEBNext Multiplex Small RNA Library Prep Set for Illumina (NEB, E7300S). For cDNA synthesis, Maxima H Minus Reverse Transcriptase (Thermo Fisher Scientific, EP0752) was used instead of the reverse transcriptase provided in the kit. Libraries were sequenced on a NovaSeq X Plus (Illumina).

### Quantification and Statistical Analysis

First, each sample (WT: three replicates; METTL1-KO1/2: three replicates) was processed using the tRAX pipeline^80^ to construct the reference of mature tRNAs and other non-coding small RNAs, pre-process raw paired-end reads to get merged and cleaned reads, map them to the reference and parse the mapping data to detect the reads derived from mature tRNAs (i.e., tRNA reads). For the reference, the 432 sequences of mature human tRNAs (i.e., hg38-mature-tRNAs) were downloaded from GtRNAdb.^81^ Second, the tRNA reads were mapped again to the mature tRNAs using BWA ver. 0.7.17-r1188.^82^ From the mapping data, read count data were generated using TIGAR2 by resolving the multi-mapped reads.^83^ Then, the anticodon-level read count data was obtained by summing the counts of tRNAs sharing the same anticodon. The differential expression analysis was performed using edgeR package ver. 3.36.0 with FDR < 0.05 as cut-off between WT and METTL1-KO1/2, respectively.^84^

### Acid urea PAGE

Acid urea PAGE was performed as previously described.^85^ Briefly, total RNAs were treated with 0.2 M Tris pH 9.5 at 42°C for 2 h. Treated and control RNA samples were then separated by acid urea PAGE.

### Fertility assay

To test male fertility, a single male fly was mated with three control (*y^1^ w^1118^*) female flies at 25°C for 3 days. After mating, the parental flies were removed, and incubation continued for 11 days (male). Emerged adult flies (progeny) were counted and the average number per vial calculated and evaluated by Tukey’s honest significant difference test. Ten independent vials were prepared for each strain. Each fertility assay was reproduced three times. The complete list of flies used in these assays is shown in Supplementary Table 3.

---

**Supplementary Figure 1.**
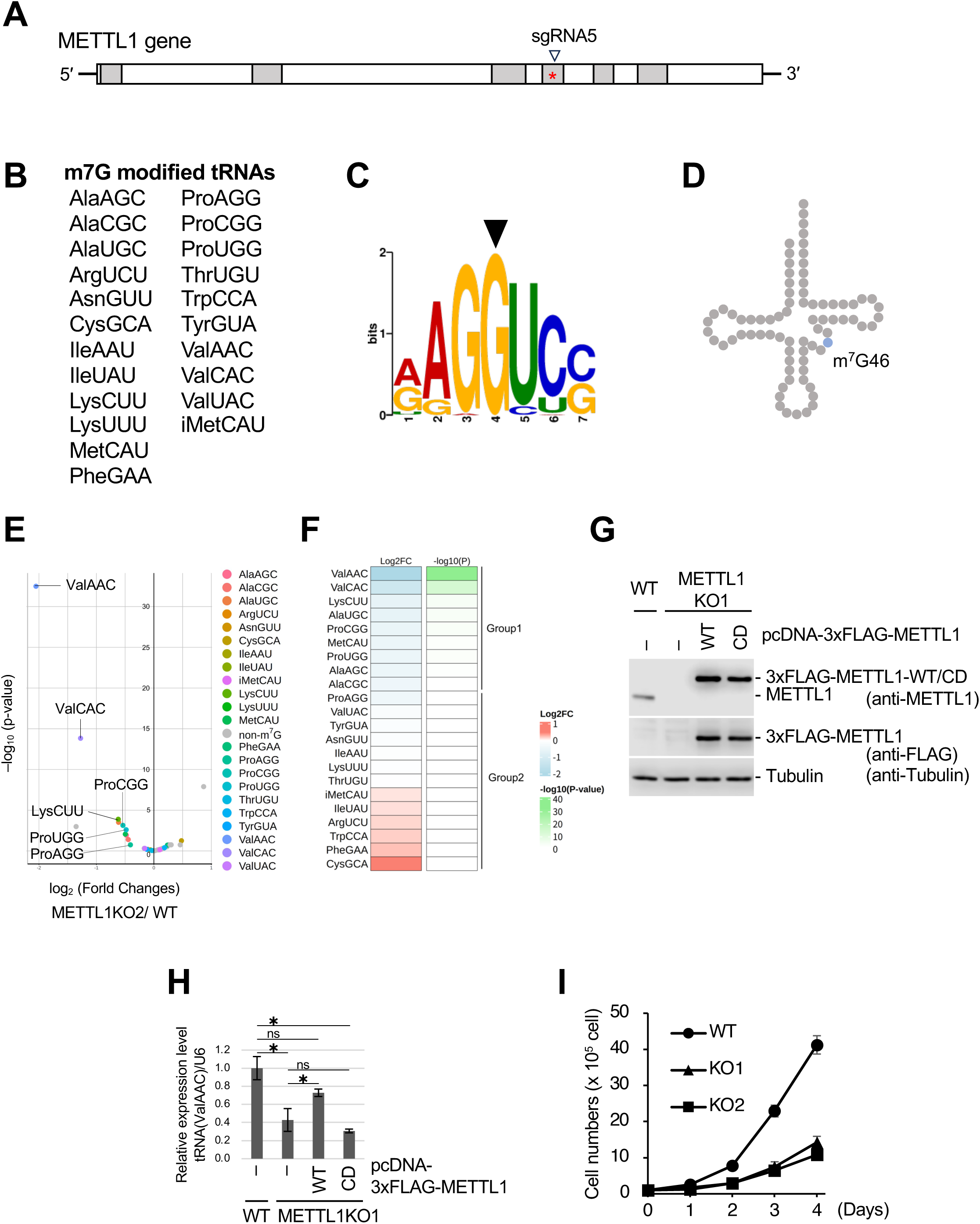
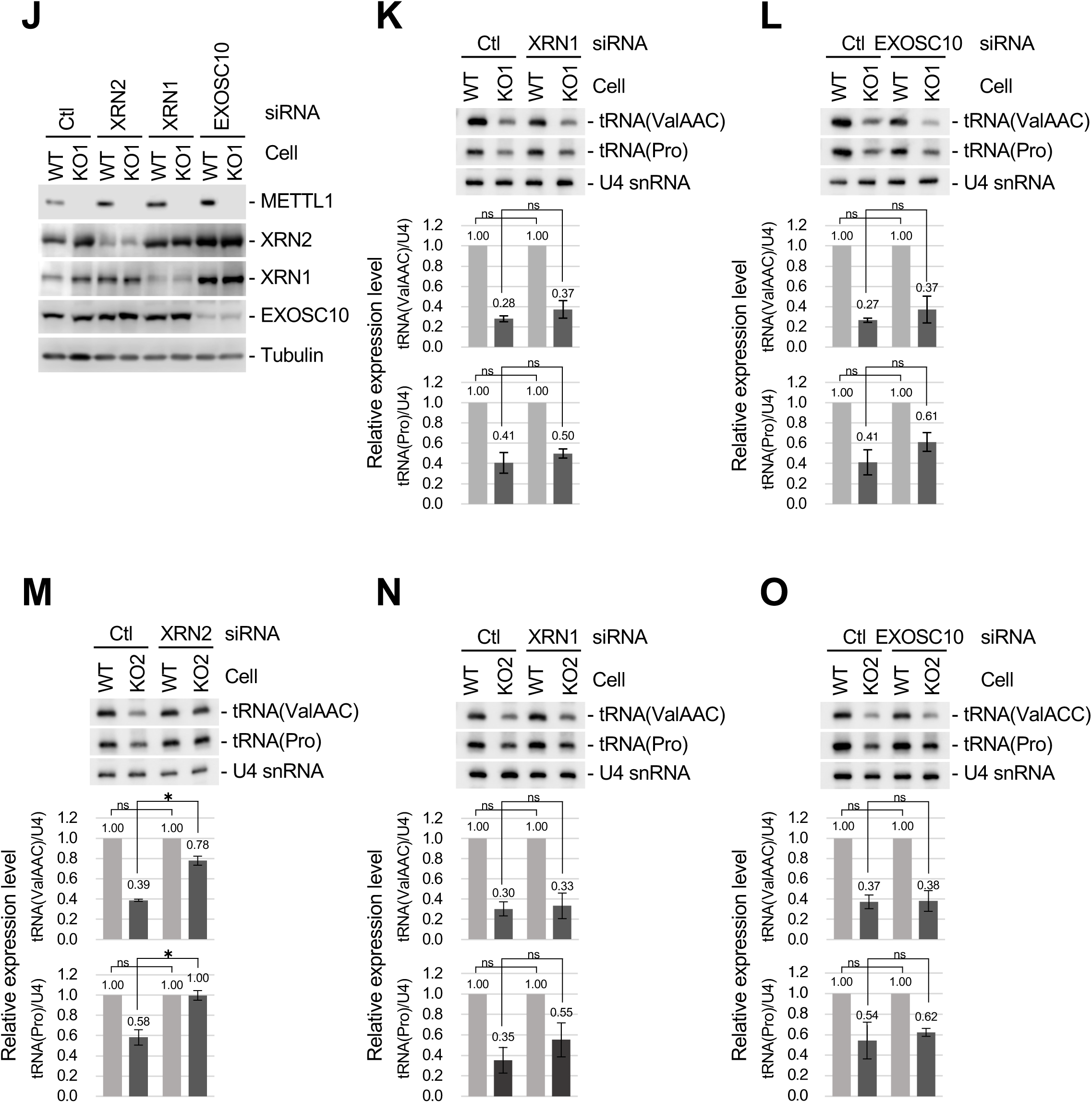
XRN2 knockdown restores m⁷G-hypomodified tRNA levels in METTL1 knockout cells (A) Schematic representation of the METTL1 gene. Gray boxes indicate exons, and the red asterisk denotes the catalytic domain responsible for the modification activity of METTL1. The arrowhead indicates the target sequences of the sgRNA. (B) List of METTL1-dependent m^7^G modified tRNAs identified in HCT116 cells by TRAC-seq. (C) Sequence motif (RAGGU motif) representing m⁷G modification sites identified in HCT116 cells by TRAC-seq. The arrowhead denotes the position of the modified guanosine (m⁷G). (D) The RAGGU motif is located in the variable loop. (E) Volcano plot of tRNA abundance changes between METTL1KO2 and WT cells. *p*-values were calculated using the Wald test with Benjamini–Hochberg correction (DESeq2 in tRAX). (F) Heatmap showing changes in m^7^G-modified tRNA abundance between METTL1KO2 and WT HCT116 cells. tRNAs were classified into three groups according to the difference in abundance between METTL1KO2 and WT, and *p*-values indicate significant differences; Group 1 (m^7^G-modified tRNAs and significantly decreased abundance, log_2_FC < 0, *p*< 0.05); Group 2 (other m^7^G-modified tRNAs, *p*≥ 0.05); Group 3 (non-m^7^G tRNAs). The *p*-values indicated correspond to those of (E). (G) Western blot showing comparable expression of 3xFLAG-METTL1 wild type (WT) and catalytic-dead mutant (CD) proteins in METTL1KO1 cells used in Figure 1E. β-Tubulin serves as a loading control. (H) Quantification of northern blot data from Figure 1E showing relative tRNA(ValAAC) abundance in WT, KO1, KO1 + METTL1-WT, and KO1 + METTL1-CD cells (mean ± s.d., n = 3). Statistical analysis was performed using one-way ANOVA (F(3, 8) = 22.5, p = 0.0003) followed by Tukey’s HSD test: WT vs KO1 (p = 0.0010), KO1 vs KO1+METTL1-WT (p = 0.0407), WT vs KO1+METTL1-WT (p = 0.1036), KO1 vs KO1+METTL1-CD (*p* = 0.3055). Bars represent mean ± s.d.; *p* < 0.05 was considered significant. (I) Cell proliferation rate of METTL1KO cells. (J) Western blot analysis confirming siRNA-mediated knockdown of XRN1, XRN2, and EXOSC10 in two independent METTL1 knockout HCT116 cell lines (KO1: METTL1KO1). Specific primary antibodies against each protein were used. (K–O) Northern blot analysis of individual tRNAs in METTL1KO1/KO2 cells transfected with the indicated siRNAs. Ctl, control siRNA. The probe for tRNA(Pro) recognizes multiple isoacceptors, including tRNA(ProAGG), tRNA(ProCGG), and tRNA(ProUGG). The lower graphs show the relative expression levels of tRNA(ValAAC) and tRNA(Pro), quantified from the northern blot signals using U4 snRNA as a loading control. Data represent mean ± s.d. from three independent experiments. Statistical significance was determined using an unpaired two-tailed Welch’s t-test (\**p* < 0.05, ns = not significant).

**Supplementary Figure 2.**
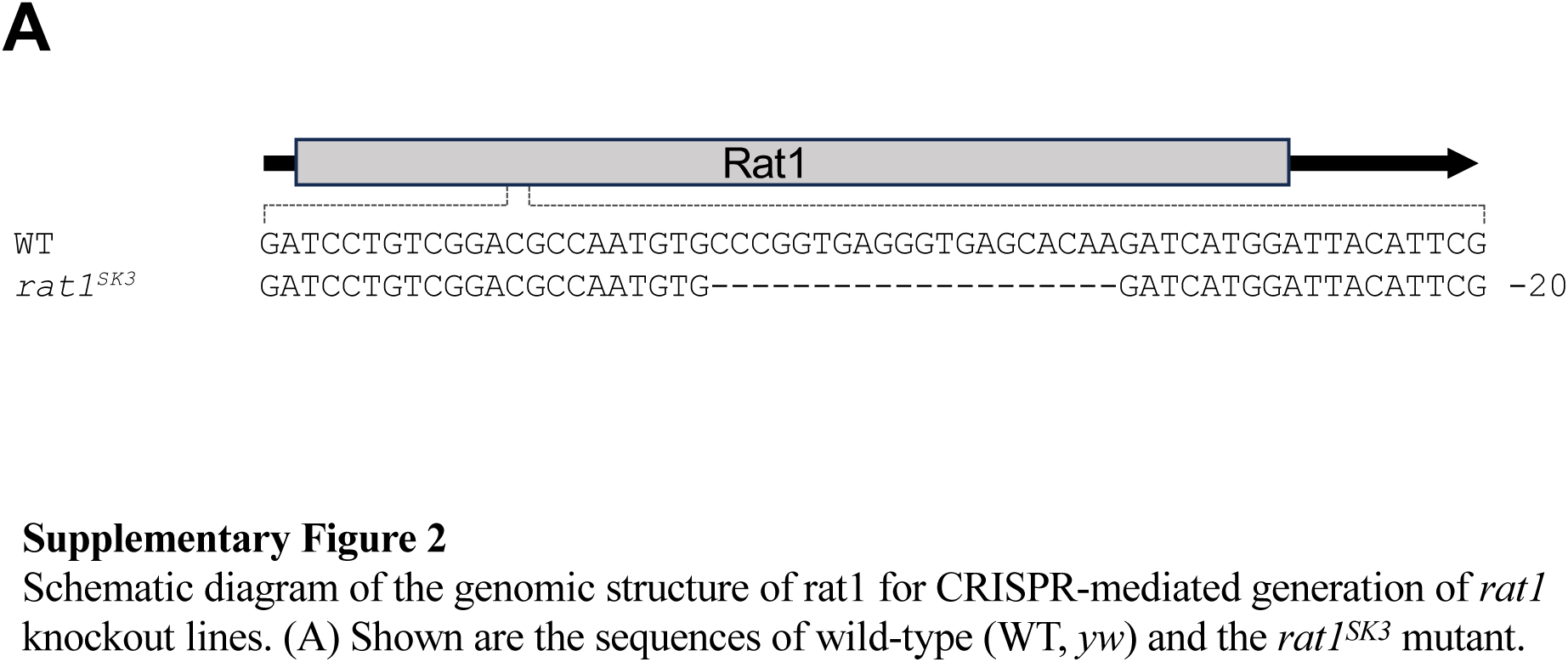
Schematic diagram of the genomic structure of rat1 for CRISPR-mediated generation of *rat1* knockout lines. (A) Shown are the sequences of wild-type (WT, *yw*) and the *rat1^SK3^* mutant.

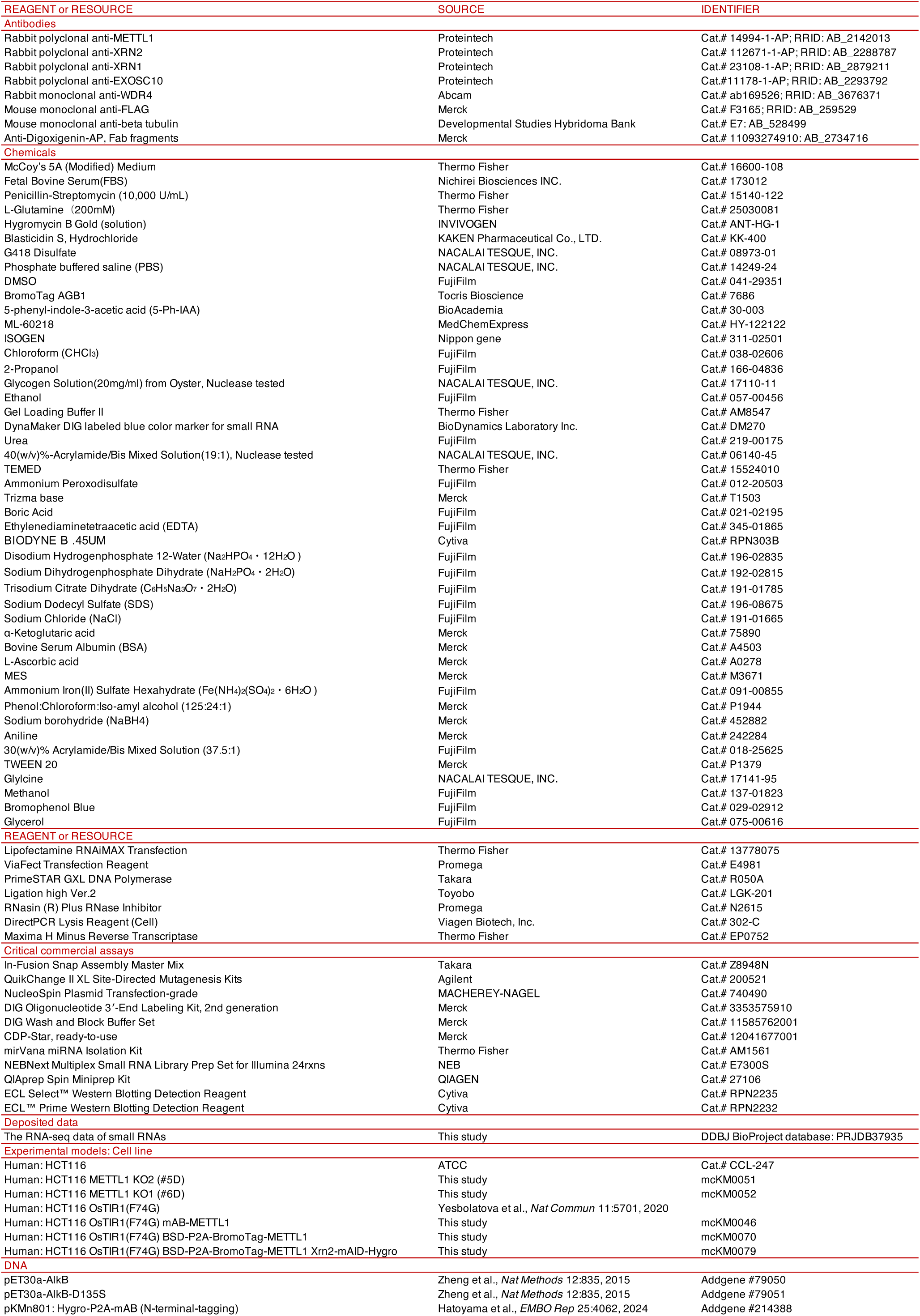
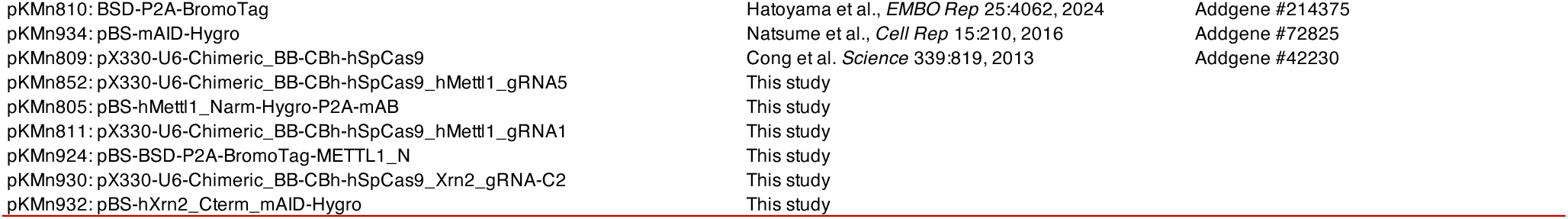

**Supplementary Table 1.**
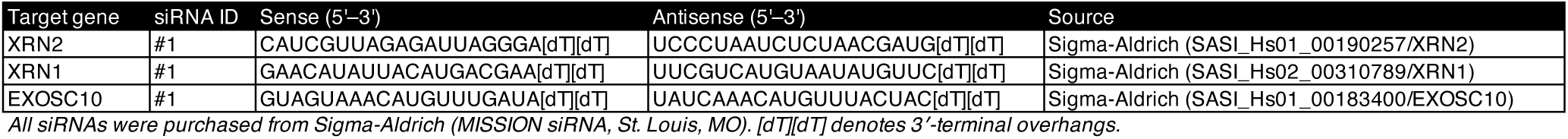
List of siRNAs used in this study.

**Supplementary Table 2.**
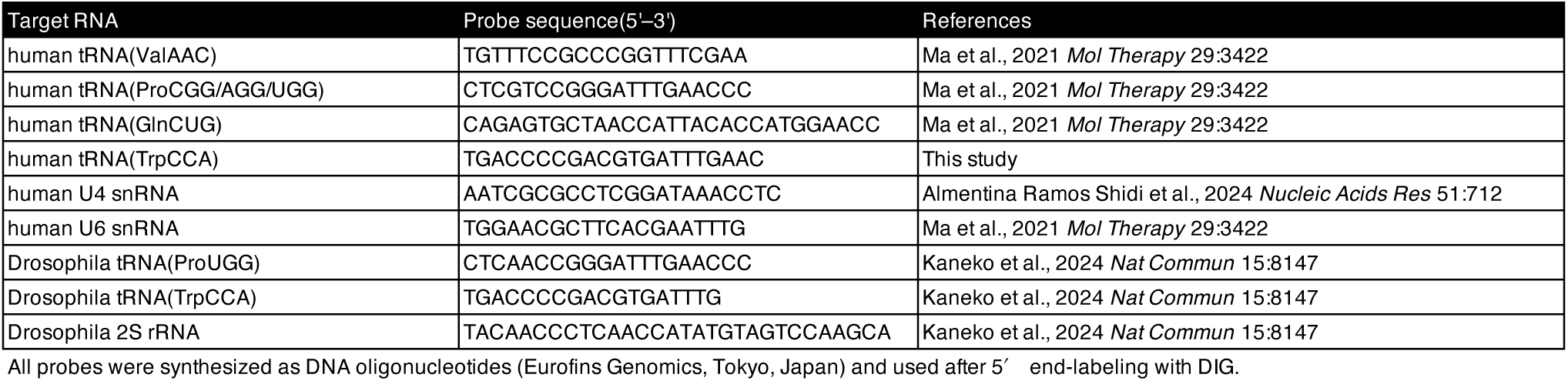
List of DNA probes used in this study.

**Supplementary Table 3.**
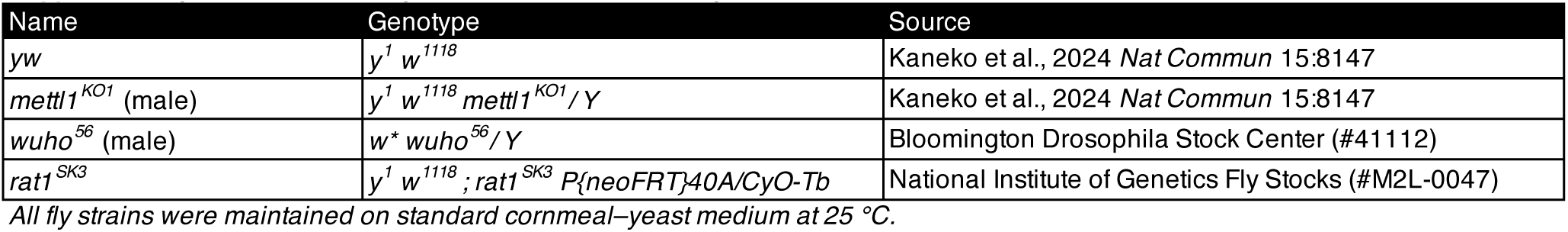
List of fly strains used in this study.

## Notes

### Competing Interest Statement

The authors have declared no competing interest.

## References

1. Cappannini, A., Ray, A., Purta, E., Mukherjee, S., Boccaletto, P., Moafinejad, S.N., Lechner, A., Barchet, C., Klaholz, B.P., Stefaniak, F., et al. (2024). MODOMICS: a database of RNA modifications and related information. 2023 update. Nucleic Acids Res. 52, D239–D244.

2. Delaunay, S., Helm, M., and Frye, M. (2024). RNA modifications in physiology and disease: towards clinical applications. Nat. Rev. Genet. 25, 104–122.

3. Hopper, A.K., and Phizicky, E.M. (2003). tRNA transfers to the limelight. Genes Dev. 17, 162–180.

4. Phizicky, E.M., and Hopper, A.K. (2010). tRNA biology charges to the front. Genes Dev. 24, 1832–1860.

5. El Yacoubi, B., Bailly, M., and de Crécy-Lagard, V. (2012). Biosynthesis and function of posttranscriptional modifications of transfer RNAs. Annu. Rev. Genet. 46, 69–95.

6. Suzuki, T. (2021). The expanding world of tRNA modifications and their disease relevance. Nat. Rev. Mol. Cell Biol. 22, 375–392.

7. Phizicky, E.M., and Hopper, A.K. (2023). The life and times of a tRNA. RNA 29, 898–957.

8. Ibba, M., and Soll, D. (2000). Aminoacyl-tRNA synthesis. Annu. Rev. Biochem. 69, 617–650.

9. Schimmel, P. (2018). The emerging complexity of the tRNA world: mammalian tRNAs beyond protein synthesis. Nat. Rev. Mol. Cell Biol. 19, 45–58.

10. Voorhees, R.M., and Ramakrishnan, V. (2013). Structural basis of the translational elongation cycle. Annu. Rev. Biochem. 82, 203–236.

11. Orellana, E.A., Siegal, E., and Gregory, R.I. (2022). tRNA dysregulation and disease. Nat. Rev. Genet. 23, 651–664.

12. Sarin, L.P., and Leidel, S.A. (2014). Modify or die?--RNA modification defects in metazoans. RNA Biol. 11, 1555–1567.

13. Tomikawa, C. (2018). 7-methylguanosine modifications in transfer RNA (tRNA). Int. J. Mol. Sci. 19, 4080.

14. Alexandrov, A., Grayhack, E.J., and Phizicky, E.M. (2005). tRNA m7G methyltransferase Trm8p/Trm82p: evidence linking activity to a growth phenotype and implicating Trm82p in maintaining levels of active Trm8p. RNA 11, 821–830.

15. Alexandrov, A., Martzen, M.R., and Phizicky, E.M. (2002). Two proteins that form a complex are required for 7-methylguanosine modification of yeast tRNA. RNA 8, 1253–1266.

16. Leulliot, N., Chaillet, M., Durand, D., Ulryck, N., Blondeau, K., and van Tilbeurgh, H. (2008). Structure of the yeast tRNA m7G methylation complex. Structure 16, 52– 61.

17. Li, J., Wang, L., Hahn, Q., Nowak, R.P., Viennet, T., Orellana, E.A., Roy Burman, S.S., Yue, H., Hunkeler, M., Fontana, P., et al. (2023). Structural basis of regulated m7G tRNA modification by METTL1-WDR4. Nature 613, 391–397.

18. Ruiz-Arroyo, V.M., Raj, R., Babu, K., Onolbaatar, O., Roberts, P.H., and Nam, Y. (2023). Structures and mechanisms of tRNA methylation by METTL1-WDR4. Nature 613, 383–390.

19. Lin, S., Liu, Q., Lelyveld, V.S., Choe, J., Szostak, J.W., and Gregory, R.I. (2018). Mettl1/Wdr4-Mediated m7G tRNA Methylome Is Required for Normal mRNA Translation and Embryonic Stem Cell Self-Renewal and Differentiation. Mol. Cell 71, 244–255.e5.

20. Alexandrov, A., Martzen, M.R., and Phizicky, E.M. (2002). Two proteins that form a complex are required for 7-methylguanosine modification of yeast tRNA. RNA 8, 1253–1266.

21. Alexandrov, A., Chernyakov, I., Gu, W., Hiley, S.L., Hughes, T.R., Grayhack, E.J., and Phizicky, E.M. (2006). Rapid tRNA decay can result from lack of nonessential modifications. Mol. Cell 21, 87–96.

22. Dewe, J.M., Whipple, J.M., Chernyakov, I., Jaramillo, L.N., and Phizicky, E.M. (2012). The yeast rapid tRNA decay pathway competes with elongation factor 1A for substrate tRNAs and acts on tRNAs lacking one or more of several modifications. RNA 18, 1886–1896.

23. Chernyakov, I., Whipple, J.M., Kotelawala, L., Grayhack, E.J., and Phizicky, E.M. (2008). Degradation of several hypomodified mature tRNA species in Saccharomyces cerevisiae is mediated by Met22 and the 5’-3’ exonucleases Rat1 and Xrn1. Genes Dev. 22, 1369–1380.

24. Kotelawala, L., Grayhack, E.J., and Phizicky, E.M. (2008). Identification of yeast tRNA Um(44) 2’-O-methyltransferase (Trm44) and demonstration of a Trm44 role in sustaining levels of specific tRNA(Ser) species. RNA 14, 158–169.

25. Whipple, J.M., Lane, E.A., Chernyakov, I., D’Silva, S., and Phizicky, E.M. (2011). The yeast rapid tRNA decay pathway primarily monitors the structural integrity of the acceptor and T-stems of mature tRNA. Genes Dev. 25, 1173–1184.

26. Guy, M.P., Young, D.L., Payea, M.J., Zhang, X., Kon, Y., Dean, K.M., Grayhack, E.J., Mathews, D.H., Fields, S., and Phizicky, E.M. (2014). Identification of the determinants of tRNA function and susceptibility to rapid tRNA decay by high-throughput in vivo analysis. Genes Dev. 28, 1721–1732.

27. Payea, M.J., Sloma, M.F., Kon, Y., Young, D.L., Guy, M.P., Zhang, X., De Zoysa, T., Fields, S., Mathews, D.H., and Phizicky, E.M. (2018). Widespread temperature sensitivity and tRNA decay due to mutations in a yeast tRNA. RNA 24, 410–422.

28. Dichtl, B., Stevens, A., and Tollervey, D. (1997). Lithium toxicity in yeast is due to the inhibition of RNA processing enzymes. EMBO J. 16, 7184–7195.

29. Yun, J.-S., Yoon, J.-H., Choi, Y.J., Son, Y.J., Kim, S., Tong, L., and Chang, J.H. (2018). Molecular mechanism for the inhibition of DXO by adenosine 3’,5’-bisphosphate. Biochem. Biophys. Res. Commun. 504, 89–95.

30. De Zoysa, T., Hauke, A.C., Iyer, N.R., Marcus, E., Ostrowski, S.M., Stegemann, F., Ermolenko, D.N., Fay, J.C., and Phizicky, E.M. (2024). A connection between the ribosome and two S. pombe tRNA modification mutants subject to rapid tRNA decay. PLoS Genet. 20, e1011146.

31. De Zoysa, T., and Phizicky, E.M. (2020). Hypomodified tRNA in evolutionarily distant yeasts can trigger rapid tRNA decay to activate the general amino acid control response, but with different consequences. PLoS Genet. 16, e1008893.

32. Kadaba, S., Krueger, A., Trice, T., Krecic, A.M., Hinnebusch, A.G., and Anderson, J. (2004). Nuclear surveillance and degradation of hypomodified initiator tRNAMet in S. cerevisiae. Genes Dev. 18, 1227–1240.

33. Kadaba, S., Wang, X., and Anderson, J.T. (2006). Nuclear RNA surveillance in Saccharomyces cerevisiae: Trf4p-dependent polyadenylation of nascent hypomethylated tRNA and an aberrant form of 5S rRNA. RNA 12, 508–521.

34. Wang, X., Jia, H., Jankowsky, E., and Anderson, J.T. (2008). Degradation of hypomodified tRNA(iMet) in vivo involves RNA-dependent ATPase activity of the DExH helicase Mtr4p. RNA 14, 107–116.

35. LaCava, J., Houseley, J., Saveanu, C., Petfalski, E., Thompson, E., Jacquier, A., and Tollervey, D. (2005). RNA degradation by the exosome is promoted by a nuclear polyadenylation complex. Cell 121, 713–724.

36. Vanácová, S., Wolf, J., Martin, G., Blank, D., Dettwiler, S., Friedlein, A., Langen, H., Keith, G., and Keller, W. (2005). A new yeast poly(A) polymerase complex involved in RNA quality control. PLoS Biol. 3, e189.

37. Tasak, M., and Phizicky, E.M. (2022). Initiator tRNA lacking 1-methyladenosine is targeted by the rapid tRNA decay pathway in evolutionarily distant yeast species. PLoS Genet. 18, e1010215.

38. Liu, N., Liu, B., Ma, C.-R., Cai, Z., Wang, J.-T., Chai, Z.-Q., Zhu, N., Shao, T., Chen, Y.-L., Lin, Y., et al. (2025). Mammalian tRNA acetylation determines translation efficiency and tRNA quality control. Nat. Commun. 16, 5496.

39. Watanabe, K., Miyagawa, R., Tomikawa, C., Mizuno, R., Takahashi, A., Hori, H., and Ijiri, K. (2013). Degradation of initiator tRNAMet by Xrn1/2 via its accumulation in the nucleus of heat-treated HeLa cells. Nucleic Acids Res. 41, 4671–4685.

40. Okamoto, M., Fujiwara, M., Hori, M., Okada, K., Yazama, F., Konishi, H., Xiao, Y., Qi, G., Shimamoto, F., Ota, T., et al. (2014). tRNA modifying enzymes, NSUN2 and METTL1, determine sensitivity to 5-fluorouracil in HeLa cells. PLoS Genet. 10, e1004639.

41. Kaneko, S., Miyoshi, K., Tomuro, K., Terauchi, M., Tanaka, R., Kondo, S., Tani, N., Ishiguro, K.-I., Toyoda, A., Kamikouchi, A., et al. (2024). Mettl1-dependent m7G tRNA modification is essential for maintaining spermatogenesis and fertility in Drosophila melanogaster. Nat. Commun. 15, 8147.

42. Braun, D.A., Shril, S., Sinha, A., Schneider, R., Tan, W., Ashraf, S., Hermle, T., Jobst-Schwan, T., Widmeier, E., Majmundar, A.J., et al. (2018). Mutations in WDR4 as a new cause of Galloway-Mowat syndrome. Am. J. Med. Genet. A 176, 2460– 2465.

43. Shaheen, R., Abdel-Salam, G.M.H., Guy, M.P., Alomar, R., Abdel-Hamid, M.S., Afifi, H.H., Ismail, S.I., Emam, B.A., Phizicky, E.M., and Alkuraya, F.S. (2015). Mutation in WDR4 impairs tRNA m(7)G46 methylation and causes a distinct form of microcephalic primordial dwarfism. Genome Biol. 16, 210.

44. Trimouille, A., Lasseaux, E., Barat, P., Deiller, C., Drunat, S., Rooryck, C., Arveiler, B., and Lacombe, D. (2018). Further delineation of the phenotype caused by biallelic variants in the WDR4 gene. Clin. Genet. 93, 374–377.

45. Hadjigeorgiou, G.M., Kountra, P.-M., Koutsis, G., Tsimourtou, V., Siokas, V., Dardioti, M., Rikos, D., Marogianni, C., Aloizou, A.-M., Karadima, G., et al. (2019). Replication study of GWAS risk loci in Greek multiple sclerosis patients. Neurol. Sci. 40, 253–260.

46. Wang, Y.-J., Mugiyanto, E., Peng, Y.-T., Huang, W.-C., Chou, W.-H., Lee, C.-C., Wang, Y.-S., Irham, L.M., Perwitasari, D.A., Hsu, M.-I., et al. (2021). Genetic association of the functional WDR4 gene in male fertility. J. Pers. Med. 11, 760.

47. Zhao, P., Xia, L., Chen, D., Xu, W., Guo, H., Xu, Y., Yan, B., Wu, X., Li, Y., Zhang, Y., et al. (2024). METTL1 mediated tRNA m7G modification promotes leukaemogenesis of AML via tRNA regulated translational control. Exp. Hematol. Oncol. 13, 8.

48. Ma, J., Han, H., Huang, Y., Yang, C., Zheng, S., Cai, T., Bi, J., Huang, X., Liu, R., Huang, L., et al. (2021). METTL1/WDR4-mediated m7G tRNA modifications and m7G codon usage promote mRNA translation and lung cancer progression. Mol. Ther. 29, 3422–3435.

49. Xia, X., Wang, Y., and Zheng, J.C. (2023). Internal m7G methylation: A novel epitranscriptomic contributor in brain development and diseases. Mol. Ther. Nucleic Acids 31, 295–308.

50. Guy, M.P., Shaw, M., Weiner, C.L., Hobson, L., Stark, Z., Rose, K., Kalscheuer, V.M., Gecz, J., and Phizicky, E.M. (2015). Defects in tRNA anticodon loop 2’-O-methylation are implicated in nonsyndromic X-linked intellectual disability due to mutations in FTSJ1. Hum. Mutat. 36, 1176–1187.

51. Nagayoshi, Y., Chujo, T., Hirata, S., Nakatsuka, H., Chen, C.-W., Takakura, M., Miyauchi, K., Ikeuchi, Y., Carlyle, B.C., Kitchen, R.R., et al. (2021). Loss of Ftsj1 perturbs codon-specific translation efficiency in the brain and is associated with X-linked intellectual disability. Sci. Adv. 7, eabf3072.

52. Arango, D., Sturgill, D., Alhusaini, N., Dillman, A.A., Sweet, T.J., Hanson, G., Hosogane, M., Sinclair, W.R., Nanan, K.K., Mandler, M.D., et al. (2018). Acetylation of cytidine in mRNA promotes translation efficiency. Cell 175, 1872–1886.e24.

53. Zhang, K., Lentini, J.M., Prevost, C.T., Hashem, M.O., Alkuraya, F.S., and Fu, D. (2020). An intellectual disability-associated missense variant in TRMT1 impairs tRNA modification and reconstitution of enzymatic activity. Hum. Mutat. 41, 600– 607.

54. Blanco, S., Dietmann, S., Flores, J.V., Hussain, S., Kutter, C., Humphreys, P., Lukk, M., Lombard, P., Treps, L., Popis, M., et al. (2014). Aberrant methylation of tRNAs links cellular stress to neuro-developmental disorders. EMBO J. 33, 2020–2039.

55. Gonskikh, Y., Tirrito, C., Bommisetti, P., Mendoza-Figueroa, M.S., Stoute, J., Kim, J., Wang, Q., Song, Y., and Liu, K.F. (2025). Spatial regulation of NSUN2-mediated tRNA m5C installation in cognitive function. Nucleic Acids Res. 53. 10.1093/nar/gkae1169.

56. Tresky, R., Miyamoto, Y., Nagayoshi, Y., Yabuki, Y., Araki, K., Takahashi, Y., Komohara, Y., Ge, H., Nishiguchi, K., Fukuda, T., et al. (2024). TRMT10A dysfunction perturbs codon translation of initiator methionine and glutamine and impairs brain functions in mice. Nucleic Acids Res. 52, 9230–9246.

57. Hatoyama, Y., Islam, M., Bond, A.G., Hayashi, K.-I., Ciulli, A., and Kanemaki, M.T. (2024). Combination of AID2 and BromoTag expands the utility of degron-based protein knockdowns. EMBO Rep. 25, 4062–4077.

58. Bond, A.G., Craigon, C., Chan, K.-H., Testa, A., Karapetsas, A., Fasimoye, R., Macartney, T., Blow, J.J., Alessi, D.R., and Ciulli, A. (2021). Development of BromoTag: A “bump-and-hole”-PROTAC system to induce potent, rapid, and selective degradation of tagged target proteins. J. Med. Chem. 64, 15477–15502.

59. Natsume, T., Kiyomitsu, T., Saga, Y., and Kanemaki, M.T. (2016). Rapid protein depletion in human cells by auxin-inducible degron tagging with short homology donors. Cell Rep. 15, 210–218.

60. Yesbolatova, A., Saito, Y., Kitamoto, N., Makino-Itou, H., Ajima, R., Nakano, R., Nakaoka, H., Fukui, K., Gamo, K., Tominari, Y., et al. (2020). The auxin-inducible degron 2 technology provides sharp degradation control in yeast, mammalian cells, and mice. Nat. Commun. 11, 5701.

61. Orellana, E.A., Liu, Q., Yankova, E., Pirouz, M., De Braekeleer, E., Zhang, W., Lim, J., Aspris, D., Sendinc, E., Garyfallos, D.A., et al. (2021). METTL1-mediated m7G modification of Arg-TCT tRNA drives oncogenic transformation. Mol. Cell 81, 3323–3338.e14.

62. Dai, Z., Liu, H., Liao, J., Huang, C., Ren, X., Zhu, W., Zhu, S., Peng, B., Li, S., Lai, J., et al. (2021). N7-Methylguanosine tRNA modification enhances oncogenic mRNA translation and promotes intrahepatic cholangiocarcinoma progression. Mol. Cell 81, 3339–3355.e8.

63. Chen, Z., Zhu, W., Zhu, S., Sun, K., Liao, J., Liu, H., Dai, Z., Han, H., Ren, X., Yang, Q., et al. (2021). METTL1 promotes hepatocarcinogenesis via m7 G tRNA modification-dependent translation control. Clin. Transl. Med. 11, e661.

64. Lin, S., Liu, Q., Jiang, Y.-Z., and Gregory, R.I. (2019). Nucleotide resolution profiling of m7G tRNA modification by TRAC-Seq. Nat. Protoc. 14, 3220–3242.

65. Fu, Y., Jiang, F., Zhang, X., Pan, Y., Xu, R., Liang, X., Wu, X., Li, X., Lin, K., Shi, R., et al. (2024). Perturbation of METTL1-mediated tRNA N7-methylguanosine modification induces senescence and aging. Nat. Commun. 15, 5713.

66. Wallrath, L.L., and Elgin, S.C. (1995). Position effect variegation in Drosophila is associated with an altered chromatin structure. Genes Dev. 9, 1263–1277.

67. Wu, J., Hou, J.H., and Hsieh, T.-S. (2006). A new Drosophila gene wh (wuho) with WD40 repeats is essential for spermatogenesis and has maximal expression in hub cells. Dev. Biol. 296, 219–230.

68. Ali, R.H., Orellana, E.A., Lee, S.H., Chae, Y.-C., Chen, Y., Clauwaert, J., Kennedy, A.L., Gutierrez, A.E., Papke, D.J., Valenzuela, M., et al. (2025). A methyltransferase-independent role for METTL1 in tRNA aminoacylation and oncogenic transformation. Mol. Cell 85, 948–961.e11.

69. Zhong, G., Chang, X., Xie, W., and Zhou, X. (2024). Targeted protein degradation: advances in drug discovery and clinical practice. Signal Transduct. Target. Ther. 9, 308.

70. Békés, M., Langley, D.R., and Crews, C.M. (2022). PROTAC targeted protein degraders: the past is prologue. Nat. Rev. Drug Discov. 21, 181–200.

71. Saito, Y., and Kanemaki, M.T. (2021). Targeted protein depletion using the auxin-inducible degron 2 (AID2) system. Curr Protoc 1, e219.

72. Cong, L., Ran, F.A., Cox, D., Lin, S., Barretto, R., Habib, N., Hsu, P.D., Wu, X., Jiang, W., Marraffini, L.A., et al. (2013). Multiplex genome engineering using CRISPR/Cas systems. Science 339, 819–823.

73. Kondo, S., and Ueda, R. (2013). Highly improved gene targeting by germline-specific Cas9 expression in Drosophila. Genetics 195, 715–721.

74. Zheng, G., Qin, Y., Clark, W.C., Dai, Q., Yi, C., He, C., Lambowitz, A.M., and Pan, T. (2015). Efficient and quantitative high-throughput tRNA sequencing. Nat. Methods 12, 835‒837.

75. Miyoshi, K., Tsukumo, H., Nagami, T., Siomi, H., and Siomi, M.C. (2005). Slicer function of Drosophila Argonautes and its involvement in RISC formation. Genes Dev. 19, 2837–2848.

76. Matsuura, J., Akichika, S., Wei, F.-Y., Suzuki, T., Yamamoto, T., Watanabe, Y., Valášek, L.S., Mukasa, A., Tomizawa, K., and Chujo, T. (2024). Human DUS1L catalyzes dihydrouridine modification at tRNA positions 16/17, and DUS1L overexpression perturbs translation. Commun. Biol. 7, 1238.

77. Saito, K., Inagaki, S., Mituyama, T., Kawamura, Y., Ono, Y., Sakota, E., Kotani, H., Asai, K., Siomi, H., and Siomi, M.C. (2009). A regulatory circuit for piwi by the large Maf gene traffic jam in Drosophila. Nature 461, 1296–1299.

78. Wintermeyer, W., and Zachau, H.G. (1975). Tertiary structure interactions of 7 - methylguanosine in yeast tRNA^Phe^ as studied by borohydride reduction. FEBS Lett. 58, 306‒309.

79. Zueva, V.S., Mankin, A.S., Bogdanov, A.A., and Baratova, L.A. (1985). Specific fragmentation of tRNA and rRNA at a 7-methylguanine residue in the presence of methylated carrier RNA. Eur. J. Biochem. 146, 679–687.

80. tRNA Analysis of eXpression (tRAX): a tool for integrating analysis of tRNAs, tRNA derived small RNAs, and tRNA modifications.

81. Chan, P.P., and Lowe, T.M. (2009). GtRNAdb: a database of transfer RNA genes detected in genomic sequence. Nucleic Acids Res. 37, D93–7.

82. Li, H., and Durbin, R. (2009). Fast and accurate short read alignment with Burrows-Wheeler transform. Bioinformatics 25, 1754–1760.

83. Nariai, N., Kojima, K., Mimori, T., Sato, Y., Kawai, Y., Yamaguchi-Kabata, Y., and Nagasaki, M. (2014). TIGAR2: sensitive and accurate estimation of transcript isoform expression with longer RNA-Seq reads. BMC Genomics 15 *Suppl 10*, S5.

84. Robinson, M.D., McCarthy, D.J., and Smyth, G.K. (2010). edgeR: a Bioconductor package for differential expression analysis of digital gene expression data. Bioinformatics 26, 139–140.

85. Köhrer, C., and Rajbhandary, U.L. (2008). The many applications of acid urea polyacrylamide gel electrophoresis to studies of tRNAs and aminoacyl-tRNA synthetases. Methods 44, 129–138.

